# Multi-echo Acquisition and Thermal Denoising Advances Precision Functional Imaging

**DOI:** 10.1101/2023.10.27.564416

**Authors:** Julia Moser, Steven M. Nelson, Sanju Koirala, Thomas J. Madison, Alyssa K. Labonte, Cristian Morales Carrasco, Eric Feczko, Lucille A. Moore, Jacob T. Lundquist, Kimberly B. Weldon, Gracie Grimsrud, Kristina Hufnagle, Weli Ahmed, Michael J. Myers, Babatunde Adeyemo, Abraham Z. Snyder, Evan M. Gordon, Nico U. F. Dosenbach, Brenden Tervo-Clemmens, Bart Larsen, Steen Moeller, Essa Yacoub, Luca Vizioli, Kamil Uğurbil, Timothy O. Laumann, Chad M. Sylvester, Damien A. Fair

**Author notes:** These authors have contributed equally to this work (Timothy O. Laumann, Chad M. Sylvester, Damien A. Fair).

## Abstract

The characterization of individual functional brain organization with Precision Functional Mapping has provided important insights in recent years in adults. However, little is known about the ontogeny of inter-individual differences in brain functional organization during human development. Precise characterization of systems organization during periods of high plasticity is likely to be essential for discoveries promoting lifelong health. Obtaining precision fMRI data during development has unique challenges that highlight the importance of establishing new methods to improve data acquisition, processing, and analysis. Here, we investigate two methods that can facilitate attaining this goal: multi-echo (ME) data acquisition and thermal noise removal with Noise Reduction with Distribution Corrected (NORDIC) principal component analysis. We applied these methods to precision fMRI data from adults, children, and newborn infants. In adults, both ME acquisitions and NORDIC increased temporal signal to noise ratio (tSNR) as well as the split-half reliability of functional connectivity matrices, with the combination helping more than either technique alone. The benefits of NORDIC denoising replicated in both our developmental samples. ME acquisitions revealed longer and more variable T2* relaxation times across the brain in infants relative to older children and adults, leading to major differences in the echo weighting for optimally combining ME data. This result suggests ME acquisitions may be a promising tool for optimizing developmental fMRI, albeit application in infants needs further investigation. The present work showcases methodological advances that improve Precision Functional Mapping in adults and developmental populations and, at the same time, highlights the need for further improvements in infant specific fMRI.

## 1. Introduction

Brain regions that are functionally connected can be identified by correlation analysis of blood oxygen level dependent (BOLD) signals -the endogenous contrast used for functional magnetic resonance imaging (fMRI; Biswal et al., 1995; Ogawa et al., 1990). Although common patterns of functional brain organization have been identified via group average studies of functional connectivity among brain regions (e.g., Gordon et al., 2016; Power et al., 2011; Yeo et al., 2011), considerable inter-individual variability exists around these global patterns (Gordon et al., 2017). Under the right conditions, these patterns are fairly stable within individuals (Gratton et al., 2018; Laumann et al., 2015). Precision Functional Mapping (PFM) is a technique that allows for a reliable characterization of such person specific functional systems (i.e., network maps or brain topography, and connection strengths or brain topology), traditionally made possible by acquiring abundant data (i.e., several hours) across multiple imaging sessions of the same participant (Gordon et al., 2017).

### Precision Functional Mapping in developmental neuroimaging

The ability to reliably and precisely map functional systems in individuals has provided an important platform for a whole host of discoveries in recent years in adults. Insights from PFM, for example, have led to a rethinking of the classic homunculus (Penfield & Boldrey, 1937) detecting inter-effector regions that interrupt effector-specific areas, forming the somato-cognitive action network (Gordon et al., 2023) and expanded our understanding of experience dependent plasticity (Newbold et al., 2020). Furthermore, recent evidence has revealed distinct patterns of expansion of the salience network in individuals with major depression (Lynch et al., 2023), demonstrating that PFM can provide important information for designing targeted neuromodulation treatments (Elbau et al., 2023). Functional brain organization in adults is as personalized as a fingerprint (Miranda-Dominguez et al., 2014) with areas of high and low probability of network overlap between persons (Hermosillo et al., 2024). Even during childhood and adolescence, personalized network topography is providing insights into the associations between brain development, cognition, age (Cui et al., 2020), and psychopathology (Cui et al., 2022). However, despite these accumulating discoveries, little is known about the ontogeny of individual differences in brain functional organization during the earliest periods of life, when individual-specific brain organization may be most informative about lifelong developmental trajectories.

The commonly used strategy of acquiring large amounts of data across multiple sessions for PFM has major limitations in many populations. For example, gathering large amounts of fMRI data in newborns and infants requires research teams to adopt specialized data acquisition strategies (Dubois et al., 2021; Korom et al., 2021) not necessarily amenable to repeated sessions. In addition, in most cases, data are acquired during natural sleep, when spontaneous waking can limit the acquisition of large amounts of, motion-free data. These data acquisition challenges are accompanied by additional inherent technical challenges of infant imaging including the lack of commercially available head coils optimized for developmental populations, resulting in sub-optimal signal to noise ratio (SNR) when using adult coils (Dubois et al., 2021). SNR can be improved by spatially smoothing or averaging data across parcels. However, these strategies lead to decreased spatial precision, which is the crux of personalized PFM. Even where large amounts of data can be collected across multiple sessions, PFM in developmental populations is still limited simply by the fast pace at which the brain develops, particularly during early stages of maturation. The rapid growth rate of the infant brain requires collecting multiple datasets in a much shorter time-period compared to a child or an adult sample for which multiple sessions of low-motion data can be collected over the course of weeks to months. Taken together, studies investigating individual differences in functional brain architecture require data that are both spatially precise and reliable to generate personalized functional brain maps, which are features that are particularly challenging to achieve in developmental samples.

### These challenges call for improvements in data collection and processing methods

These challenges highlight the importance of identifying data acquisition, processing, and analysis strategies that maximize signal and reduce the time in the scanner needed to obtain reliable characterization of functional network topographies and their underlying topology. This paper examines two possible methodological improvements for developmental neuroimaging that previously and independently have been shown to increase signal quality and reliability in adult participants: 1) Multi-echo (ME) data acquisition (Lynch et al., 2020) and 2) Noise Reduction with Distribution Corrected (NORDIC) principal component analysis (Dowdle et al., 2023; Moeller et al., 2021; Vizioli et al., 2021).

In contrast to traditional BOLD imaging, ME data acquisitions capture images at multiple echo times during a single readout time of the T2*, or transverse relaxation signal decay. The T2* weighted fMRI signal reflects the decay in transverse magnetization introduced by the radio frequency (RF) pulse. Following the RF pulse, the fMRI signal decays exponentially over successive echoes (Posse, (2012); see Figure 5A for an example). T2* relaxation times vary across age, brain regions and tissue types, due to differences in underlying neurobiological tissue properties. The optimal echo time to capture an image (i.e., tradeoff between signal to noise and functional contrast) is usually defaulted to the T2* relaxation time of the voxels of interest. Voxels with longer T2*s have higher signal intensities in images from longer echo times compared to voxels with shorter T2*s (Kundu et al., 2017). With ME data acquisition, data from different echo times can be optimally combined based on the T2* of the underlying tissue (Posse et al., 1999). The properties of the signal decay across echoes can furthermore be used to separate BOLD effects from non-BOLD effects using multi-echo independent component analysis (MEICA; Kundu et al., (2012)).

The technique of ME data acquisition has been known for some time (e.g., Posse et al., (1999)) but has struggled to gain popularity owing to the requisite compromises in spatial and temporal resolution as well as the need to use in-plane accelerations to avoid excessively long echo trains during image readout. NORDIC, a recently developed tool for denoising fMRI data, could help to overcome some of these challenges and further improve the capabilities and usability of ME data. NORDIC enhances image quality by removing zero-mean, unstructured thermal noise, thereby improving the SNR without sacrificing spatial precision (Dowdle et al., 2023; Vizioli et al., 2021). However, little is known about the value of ME-NORDIC fMRI, particularly in pediatric populations.

### Translating methods from adults to infants - challenges and opportunities

Most methodological advances are first established in healthy young adults. However, their transferability to developmental neuroimaging is not guaranteed. Particularly for very young populations, it is important to consider that infant brains are not merely smaller adult brains but instead have specific properties that change with their developmental stage. T2* relaxation, which forms the basis of fMRI, shows a developmental trend, being slower in newborns compared to infants and adults (Rivkin et al., 2004; Williams et al., 2005). This slower T2* decay is related to several factors, including reduction in the amount of free water compared to bound water in brain tissues over development (Engelbrecht et al., 1998; Xydis et al., 2006) as axon myelination increases in both white and gray matter (Baumann & Pham-Dinh, 2001; Dubois et al., 2014; Kostović et al., 2019). Other factors are an increase in proton density, an abundance of macromolecules, and changes in iron concentration (Goksan et al., 2017). These are important factors to consider for the evaluation of a prospective ME acquisition in this age group as the optimal combination of echoes is dependent on the T2* relaxation time of a given voxel. Given the impact of brain developmental changes on T2* relaxation times, ME data acquisitions could indeed be a promising tool for developmental imaging. It opens up the possibility to account for the impact of brain developmental differences on fMRI signals. This may have particular benefit for longitudinal investigations, in which individually optimized echo combinations based on T2* times could be determined for each acquisition time point while still using the same acquisition sequence.

### Methods under investigation in the present study

The present study is an individualized study in which we investigated T2* relaxation times and optimal echo combinations of ME fMRI data across different ages and the impact of NORDIC thermal noise removal on these data. To test the usability and benefit of these methodological approaches, we utilized precision imaging data with single-echo (SE) and ME data acquisitions from adults, children, and newborns acquired across multiple days. We compared ME infant and adult data acquired with the same sequence to gain a better understanding of ME data acquisition and optimal echo times in infants compared to adults, for whom ME fMRI previously been shown to provide advantages over SE acquisitions (Lynch et al., 2020). MEICA is not included in the present investigation as its ability to separate BOLD effects from non-BOLD effects is based on signal decay curve shapes, which are different in young infants and not sufficiently understood yet. We used NORDIC denoising on all datasets to look at differences in data quality with and without thermal noise removal. We hypothesize that the application of NORDIC denoising with its targeting of thermal noise, will be useful independent of the age of the target sample. Data quality was evaluated using temporal SNR (tSNR), strength of functional connections, and split-half reliability of brain functional connectivity within individuals. These metrics will allow us to see if effects of NORDIC and ME optimal echo combination on tSNR are additive or multiplicative or redundant. This investigation informs the application of these methods to developmental neuroimaging with the goal of improving signal quality to facilitate precision functional imaging.

## 2. Methods

### 2.1 Sample

The sample consists of one adult (PA001) and two children (PC001 & PC002; age 10) enrolled at the University of Minnesota (UMN), and one adult (PA002) and three healthy neonates enrolled at Washington University in St. Louis (WashU), ages 28 days (43 weeks postmenstrual age (wPMA); PB004), 12 days (41 wPMA; PB005) and 13 days (41 wPMA; PB001). Data included in this manuscript were collected for studies approved by the UMN’s Institutional Review Board and the Human Studies Committees at WashU, and written informed consent was obtained from all participants and parents of minor participants. For PA001, ME and SE data were acquired in four sessions over three months. For PA002 ME data were acquired in six sessions over four months and SE data were acquired in three sessions over one month. For PC001 and PC002 data were acquired in four sessions over eight months, and for neonatal participants, data were acquired over four to five days within one week. Anatomical and resting state data in neonates were acquired during natural sleep.

### 2.2 Data acquisition

rs-fMRI data for all subjects was acquired with the CMRR multiband (MB)-ME sequence (Feinberg et al., 2010; Moeller et al., 2010) with 5 echoes at WashU and 4 echoes at UMN - dropping the last echo based on compatibility issues with Siemens XA30 (14.2ms, 38.93ms, 63.66ms, 88.39ms, 113.12ms; TR = 1.761s, 2mm resolution, MB factor = 6, IPAT = 2, flip angle = 68°). Data were acquired in PA phase encoding direction. The total amount of rs-fMRI data acquired ranged between 72 minutes (PB004) and 203 minutes (PA002; see Suppl. Table 1). Additionally, spin echo fieldmaps were acquired in both AP and PA direction (SE, single-band, 3 frames per run, TR = 8.0s, TE = 66ms, flip angle = 90°) for all participants.

We additionally acquired single-echo rs-fMRI data for PA001, PA002, PB004 and PB005 with a more standard fMRI acquisition protocol (TR 0.8s, TE = 37ms, 2mm resolution, MB factor = 8, flip angle = 52°; infants: TR 1.51s, TE = 37ms, 2mm resolution, MB factor = 4, flip angle = 52°, PA phase encoding direction). The total amount of SE rs-fMRI data acquired ranged between 77 minutes (PB004) and 159 minutes (PA001; see Suppl. Table 1).

Imaging for PA001 and PC subjects was performed at UMN using a Siemens 3-T Prisma scanner and a 64 channel (PC001) and a 32 channel (PA001, PC002) head coil. At WashU, functional imaging for PA002 was performed using a Siemens 3-T Prisma scanner and a 64 channel head coil. Neonatal imaging was performed using a Siemens 3-T Prisma scanner and a 32 channel head coil. For details on anatomical references, see Supplemental Methods.

### 2.3 Data processing

Data for all processing and analysis was converted to BIDS format with dcm2bids (Boré et al., 2023). For analyses in which NORDIC was applied to a dataset, this step was performed before the regular preprocessing using the phase and the magnitude images of the scan data. Using phase and magnitude images for NORDIC allows to maintain complex-valued Gaussian noise (see Moeller et al., (2021) for details). Three noise frames acquired at the end of each functional run were used to help estimate the empirical thermal noise level. For the one participant in which noise scans were not acquired (PB001), we used a theoretical thermal noise level (implemented as 1/sqrt(2); Moeller et al., (2021); Vizioli et al., (2021)). NORDIC was implemented in Matlab R2019a and applied to data from each run (Vizioli et al., 2021). In the case of ME data acquisition, NORDIC denoising was performed for each echo individually.

Adult and children’s imaging data were processed using a recently upgraded version of fMRIprep (Esteban et al., 2019); version 24.0.0, development version dev55+g4d21c37a; github commit 4d21c37a), now consistent with processes outlined for ABCD-BIDS (Sturgeon, Earl, et al., 2023). Susceptibility distortion correction of BOLD time series was done within fMRIPrep, using an FSL topup-based method to estimate fieldmaps from “PEPolar” acquisitions (acquisitions with opposite phase encoding direction) (Andersson et al., 2003). These fieldmaps were then used to correct distortion of the BOLD time series data. Additional options enabled for fMRIPrep processing were “--project-goodvoxels”, to exclude voxels with locally high coefficient of variation from volume-to-surface projection of the BOLD time series, and “--cifti-output 91k” to enable output of BOLD time series and morphometric data (surface curvature, sulcal depth, and cortical thickness maps) in the HCP grayordinates space (Glasser et al., 2013). This updated version of fMRIprep optimally combines ME data within its processing workflow using T2* based echo weighting 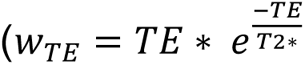; Posse et al., (1999)).

After fMRIPrep processing, functional connectivity postprocessing was performed using XCP-D 0.6.1 (Mehta et al., 2023) with options “--cifti” (ingress CIFTI fMRIPrep CIFTI derivatives), “--warp-surfaces-native2std” (apply the T1w-atlas nonlinear transform from the fMRIPrep derivatives to align XCP-D’s surface mesh output to the MNI152NLin6Asym template), “-m” (after processing, concatenate BOLD timeseries and motion data across runs), “--dcan-qc” (generate a QC report in similar format to the “executive summary” of the DCAN ABCD-BIDS pipeline), “--despike” (apply AFNI 3dDespike to BOLD timeseries data), “--lower-bpf” 0.009 (set lower bound of band-pass filter to 0.009 Hz, as in the ABCD-BIDS pipeline, the default of 0.08 is used for the “--upper-bpf’), and “–band-stop-min” <min> “–band-stop-max” <max> (set the frequency range used for respiratory motion filtering: PA001: 12-18 breaths per minute (bpm) , PA002: 11-17 bpm, PC001&2: 15-25 bpm) as well as grayordinate based signal regression. Frequency ranges for the band-stop filter were chosen based on age specific respiration rates and individual peaks in respiratory power plots (PA002). Functional data was additionally denoised and motion censored using framewise displacement (FD) with a threshold of 0.2mm.

Neonatal images were preprocessed with the infant-abcd-hcp-pipeline (Sturgeon, Snider, et al., 2023), an infant specific modification of the HCP pipeline (Feczko et al., 2021; Glasser et al., 2013). Segmentations of anatomical images were created using BIBSnet, a deep learning tool specifically trained for infant MRI image segmentation. (Hendrickson et al., 2023). These precomputed segmentations were utilized by the infant-abcd-hcp-pipeline during pre-processing. Anatomical images were registered to the MNI infant atlas and functional data was projected onto the atlas space surfaces. Spin echo fieldmaps were leveraged for susceptibility distortion correction using FSL topup (Andersson et al., 2003). Functional connectivity processing was performed during the DCANBOLDProcessing stage of the infant-abcd-bids-pipeline, which reflects the processing steps performed by XCP-D (interpolation of high-motion volumes, detrending, confound regression, bandpass filtering). Compared to adults and children, no respiratory filter was applied for the infant data as their respiration rate (30-60 bpm) is higher than half of the Nyquist frequency based on the TR. Functional data was additionally denoised and motion censored using framewise displacement (FD) with a threshold of 0.3mm. FD traces were visually inspected as recommended by (Power et al., 2012) and it was determined that 0.3 was the noise floor for infants. Runs with less than 30% of remaining data at an FD of 0.3mm were excluded. After censoring frames with high motion (above FD 0.3mm) during data preprocessing, PB004 retained 57 minutes of low motion data (out of 77), PB005 79 minutes (out of 117) and PB001 112 minutes (out of 142) with one entire run excluded. From the additional SE data, PB004 retained 72 minutes (out of 77) and PB005 retained 85 minutes (out of 123; Suppl. Table 1).

As the infant-abcd-hcp-pipeline is not constructed for ME data, data from all five echoes were optimally combined using Tedana (DuPre et al., 2021; Kundu et al., 2012, 2013; The tedana Community et al., 2022) before running the pipeline. For the optimal combination we used the T2* based echo weighting implemented in Tedana, which is also implemented in fMRIprep. In addition, motion regressors were calculated from the first echo and applied to all individual echoes before combining them.

### 2.4 Data analysis

We calculated tSNR, defined as mean intensity over the timeseries divided by the time series standard deviation, in the data with and without NORDIC applied. Averages across runs with >90% low motion data were used for tSNR estimates. This is done to ensure that this metric is not impacted by subjects’ motion while keeping the same amount of frames for each run (avoiding motion censoring). Data smoothness was estimated from preprocessed BOLD runs using 3dFWHMx (options -combine -detrend -automask -acf) in AFNI (version 16.1.13; Cox, (1996). Functional connection strength was computed using a parcel-to-parcel connectivity matrix (parcels for adults and children from Gordon et al., (2016), for neonates from Myers et al., (2024)).

Reliability was quantified as the grayordinate-by-grayordinate correlation of dense functional connectivity matrices within subject similar to (Lynch et al., 2020). Dense connectivity matrices were computed without applying any additional spatial smoothing. For constructing reliability curves, data were split in half and consecutive amounts of data compared to the held-out half. To account for natural variations in data quality during one session and across multiple days, the order of runs was randomly permuted 100 times and an average curve calculated. Reliability curves for parcellated data were calculated in the same fashion. The reliability values reported throughout the paper represent the average value of the spatial reliability map for all grayordinates or parcels.

We investigated optimal echo times (T2*) as outputted by Tedana as well as the weights of each echo used for the T2* based optimal echo combination for each grayordinate. As for tSNR estimates, T2* was averaged across runs with >90% low motion data as T2* maps are estimated during data pre-processing before motion censoring.

## 3. Results

### ME and NORDIC improved reliability while keeping spatial precision

Results from SE and ME data acquisitions in two adult precision imaging participants replicated the recently published (Lynch et al., 2020) increase in tSNR and reliability for ME. Removal of thermal noise with NORDIC furthermore increased tSNR for both SE and ME data. The highest tSNR was obtained with the combination of ME and NORDIC (Figure 1, Table 1, Suppl. Fig 1A). To measure data reliability, the available data for each participant was split in half. The connectivity matrices computed from increasing amounts of consecutive data from one half were correlated to the matrix from the held-out half. Both SE and ME acquisitions had improved reliability with NORDIC while still retaining the person’s specific connectivity structure (Figure 2, Table 2, Suppl. Fig 1B). The reliability that can be reached in PA001 with 70 minutes of SE data (M=0.24) was reached with 10-15 minutes of data with ME-NORDIC (Figure 2B). The advantage of ME was particularly evident when comparing connectivity patterns between SE and ME data in areas with high signal dropout (Figure 2C). NORDIC increased absolute connectivity strength generally across functional connections (Suppl. Fig 2, Suppl. Table 2). Importantly, loss in spatial precision for data denoised with NORDIC was minimal (FWHM increased from 2.73 to 2.8 for SE-NORDIC and from 3.47 to to 3.7 for ME-NORDIC; Suppl. Table 3). Relative to SE data, using ME and NORDIC increased FWHM by 0.96, which is minimal relative to commonly used data smoothing approaches (FWHM increased by 3.59 for Kernel 1.5 (FWHM = 6.32) and 7.95 for Kernel 2.5 (FWHM = 10.68); Suppl. Fig 3). These results demonstrate an increase in data reliability with minimal loss in spatial precision when combining ME acquisitions with NORDIC.

**Figure 1:**
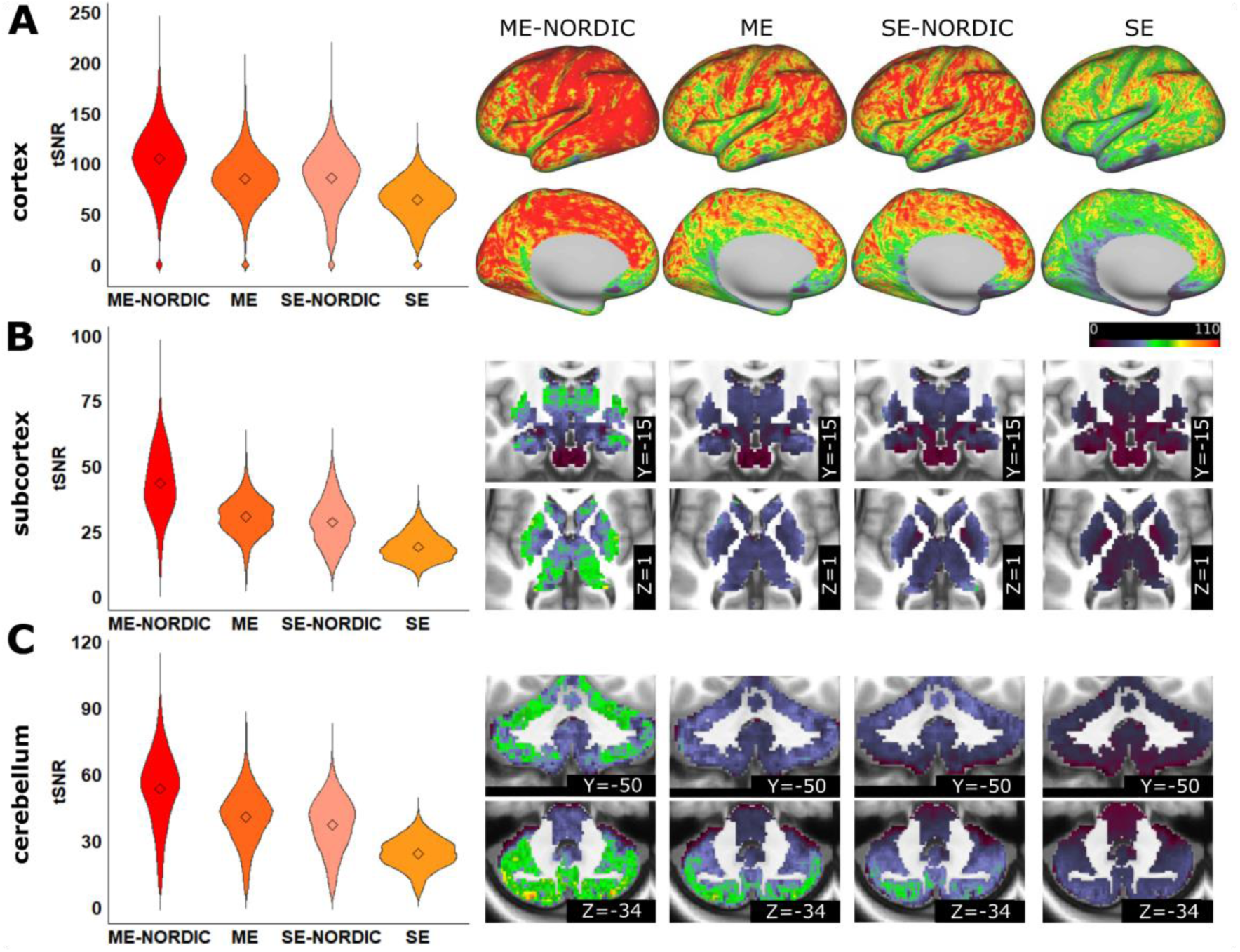
tSNR values for cortical and subcortical structures for SE and ME data with and without NORDIC for adult subject PA001 (average of runs with >90% low motion). A) Cortex, B) Subcortex, C) Cerebellum

**Figure 2:**
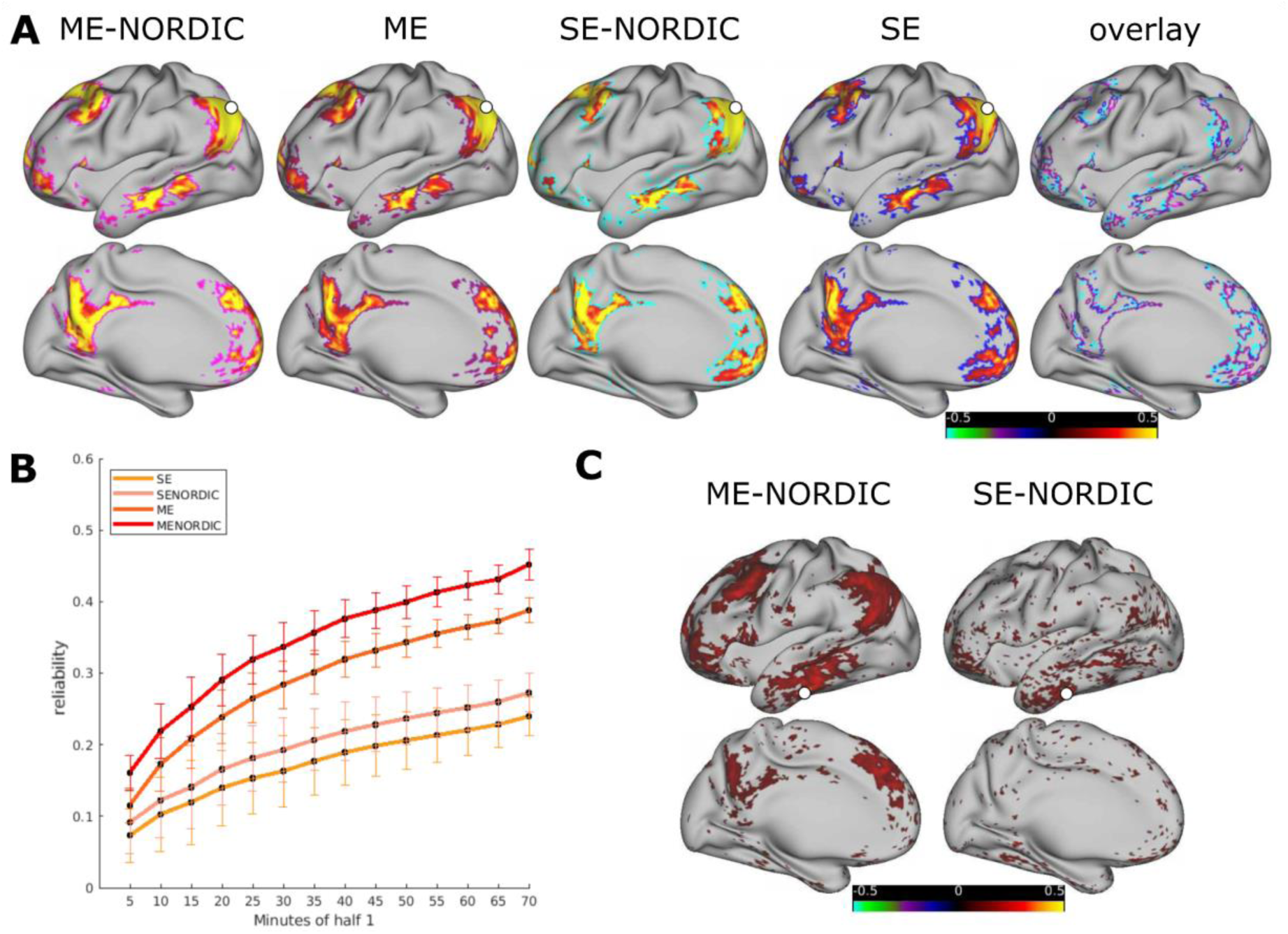
A) A person’s specific connectivity pattern (adult PA001) remains across SE and ME acquisitions with and without NORDIC. Seed based connectivity for all four data acquisition conditions (default mode network seed displayed in white). Outlined threshold shows areas of highest connectivity (top 10%). Rightmost panel shows the similarity of the spatial extent of high connectivity between conditions. B) Reliability curves for a split half of the data of PA001. Curves represent the average reliability across all grayordinates and 100 permutations of the run order. Error bars show SD across permutations. C) ME acquisitions improve signal particularly in areas with high signal dropout which helps to detect connectivity patterns in these areas. Displayed example: seed based connectivity of inferior temporal seed (seed region displayed in white) comparing ME-NORDIC and SE-NORDIC.

**Table 1:** tSNR with and without NORDIC for precision imaging participants. SD represents SD across brain grayordinates.

|  |  |  | non-NORDIC | NORDIC | Percent difference |
| --- | --- | --- | --- | --- | --- |
| adults | PA001 | ME | M = 67.93, SD = 31.11 | M = 85.11, SD = 37.32 | 25.29% |
|  |  | SE | M = 49.23, SD = 26.13 | M = 67.52, SD = 34.62 | 37.15% |
|  | PA002 | ME | M = 54.26, SD = 22.09 | M = 63.34, SD = 24.83 | 16.73% |
|  |  | SE | M = 44.13, SD = 24.25 | M = 55.09, SD = 30.11 | 24.84% |
| children | PC001 | ME | M = 67.29, SD = 29.69 | M = 79.37, SD = 32.55 | 17.95% |
|  | PC002 | ME | M = 72.27, SD = 31.84 | M = 84.06, SD = 35.84 | 16.31% |
| infants | PB004 | ME | M = 95.40, SD = 38.33 | M = 127.66, SD = 48.90 | 33.82% |
|  |  | SE | M = 90.2, SD = 38 | M = 135.56, SD = 67.63 | 50.29% |
|  | PB005 | ME | M = 68.92, SD = 27.5 | M = 81.35, SD = 30.52 | 18.04% |
|  |  | SE | M = 68.24, SD = 26.94 | M = 83.9, SD = 36.34 | 22.95% |
|  | PB001 | ME | M = 94.21, SD = 39.33 | M = 145.84, SD = 53.62 | 54.8% |

**Table 2:** Split half reliability with and without NORDIC for precision imaging participants. SD represents SD across permutations.

|  |  | minutes |  | non-NORDIC | NORDIC | Percent difference |
| --- | --- | --- | --- | --- | --- | --- |
| adults | PA001 | 70 | ME | M = 0.39, SD = 0.24 | M = 0.45, SD = 0.23 | 16.41% |
|  |  |  | SE | M = 0.24, SD = 0.21 | M = 0.27, SD = 0.23 | 13.77% |
|  | PA002 | 70 | ME | M = 0.29, SD = 0.21 | M = 0.33, SD = 0.22 | 14.19% |
|  |  |  | SE | M = 0.2, SD = 0.19 | M = 0.23, SD = 0.21 | 15.22% |
| children | PC001 | 80 | ME | M = 0.39, SD = 0.26 | M = 0.45, SD = 0.27 | 16.74% |
|  | PC002 | 70 | ME | M = 0.44, SD = 0.23 | M = 0.5, SD = 0.23 | 13.67% |
| infants | PB004 | 35 | ME | M = 0.04, SD = 0.05 | M = 0.06, SD = 0.07 | 54.47% |
|  |  |  | SE | M = 0.02, SD = 0.03 | M = 0.09, SD = 0.09 | 301.38% |
|  | PB005 | 25 | ME | M = 0.13, SD = 0.13 | M = 0.16, SD = 0.15 | 25.7% |
|  |  |  | SE | M = 0.07, SD = 0.08 | M = 0.15, SD = 1.15 | 128.09% |
|  | PB001 | 50 | ME | M = 0.11, SD = 0.12 | M = 0.17, SD = 0.15 | 51.81% |

### NORDIC improves tSNR and reliability of ME data in developmental samples

ME precision imaging data from two children and three infants were used to assess the translatability of ME-NORDIC benefits to developmental populations, in whom acquisition of large amounts of resting-state data to achieve reliable individual network solutions is less feasible. As in adults, we investigated changes in tSNR, split-half reliability, and strength of functional connections.

As in adults, we calculated the average tSNR from runs with greater than 90% of low motion data (Suppl. Table 1). We saw an overall improvement in tSNR with NORDIC for all populations (Figure 3A and Suppl. Figures 4A, 5A, & 6A Table 1). Similarly to adults, split half reliability increased with NORDIC both in children and infants (Figure 3B, Table 2, Suppl. Figures 4B, 5B, & 6B).

**Figure 3:**
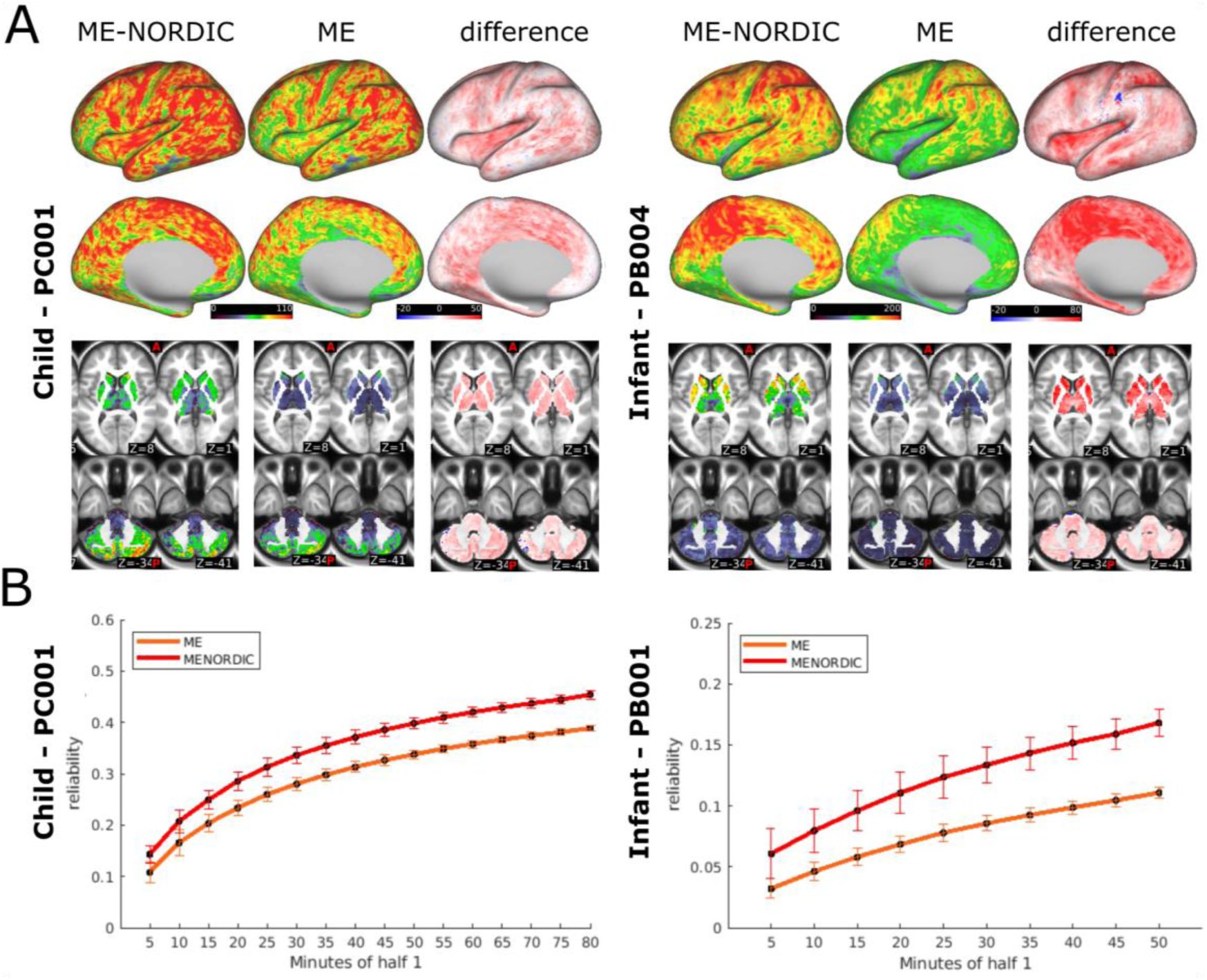
A) tSNR values for cortical and subcortical structures with and without NORDIC in child participant PC001 and infant participant PB004 (average of runs with >90% low motion). B) Gains in reliability with NORDIC for PC001 and PB001 (infant with largest amount of low motion ME data). Curves represent the average reliability across all grayordinates and 100 permutations of the run order. Error bars show SD across permutations.

Furthermore, across all participants and age groups, data denoised with NORDIC showed stronger functional connectivity (Suppl. Table 2, Suppl. Figures 7 & 8). For this comparison, we looked at parcellated connectivity matrices using the same amount of low motion data for each condition, using an age specific parcellation scheme for the infants. Similar benefits were also seen in single echo data in infants (Suppl. Figures 4, 5, & 7).

### ME acquisitions in infants provide additional insights regarding brain development

In addition to boosting reliability, ME acquisitions provide insight into fMRI signal properties. The ME data acquisition protocol allowed us to model the T2 decay curve across four/five different time points and estimate T2* relaxation times (Suppl. Figure 9 shows consistency of these estimates with and without the fifth echo). The mean T2* relaxation time across surface vertices for the adults was 48.89ms for PA002 (SD = 12.23ms, 5th and 95th percentile = [21.26ms, 64.75ms]) and 50.62ms for PA001 (SD = 12.7ms, 5th and 95th percentile = [22.23ms, 65.51ms]), which is consistent with the adult literature (Fera et al., 2004; Wansapura et al., 1999). In children, the mean T2* relaxation time was 59.34ms for PC001 (SD = 13.96ms, 5th and 95th percentile = [27.91ms, 75.18ms]) and 58.16ms for PC002 (SD = 14.08ms, 5th and 95th percentile = [24.59ms, 72.64ms]), showing a slight increase compared to adults.

T2* relaxation times were longer and more variable in all infants compared to adult or child subjects (Figure 4, Suppl. Figure 10). PB004 showed a mean T2* relaxation time of 93.84ms (SD = 27.04ms, 5th and 95th percentile = [37.23ms, 130.22ms]), PB005 77.49ms (SD = 30.46, 5th and 95th percentile = [27.43ms, 124.6ms]) and PB001 81.51ms (SD = 27.84, 5th and 95th percentile = [31.91ms, 124.10ms]). Application of NORDIC denoising to the data had a minimal and non-systematic impact on the estimate of T2* (Suppl. Figure 10).

**Figure 4:**
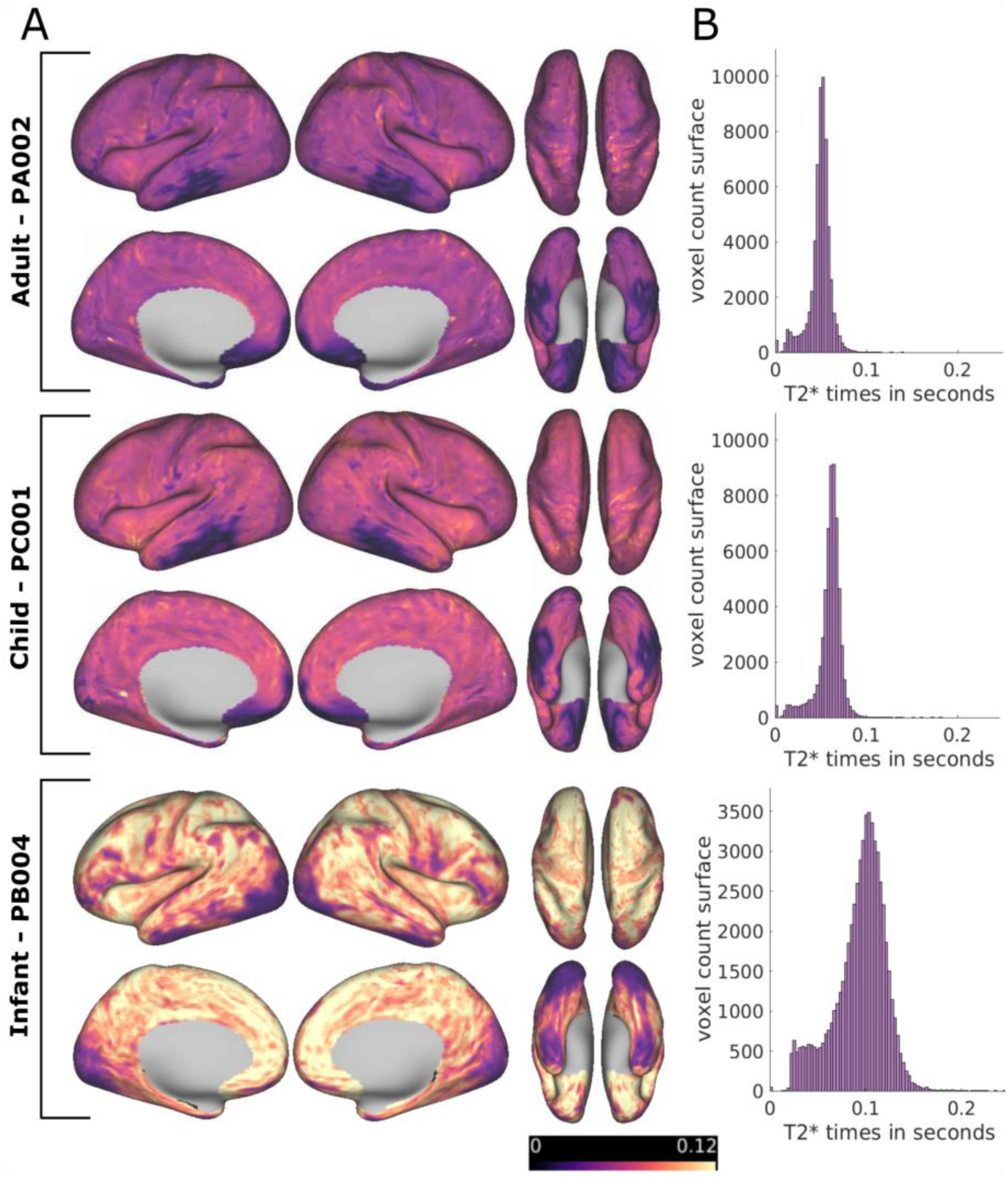
A: T2* values (in seconds) for PA002, PC001 and PB004 (average of runs with >90% low motion) computed from ME data without NORDIC denoising. B: distribution of T2* times from the cortical surface.

### Tissue properties in infants affect the optimal echo combination

The large difference in T2* relaxation times between infants and adults or children (Figure 4) highlight an important challenge for methodological advancements for developmental neuroimaging, namely the translatability of findings from adults to brains with fundamentally different properties. In this specific example, the striking difference in T2* relaxation times during early development caused an alteration in echo weighting for the optimally combined ME signal between the different age groups (Figure 5, Suppl. Figure 11). Differences in optimally combining ME acquisitions between adult subject PA002 (five-echo sequence) and the infant subjects are summarized in Table 3. The mean normalized weighting per echo in PA002 was highest for the second and third echo while, for the infants, the third to fifth echoes were weighted heaviest. In all subjects, weighting of the first echo was highest in limited areas which is likely related to signal dropout in these areas (Figure 5). The differences in echo weighting in different age groups affect tSNR, and as a corollary, the impact of NORDIC, as thermal noise contribution relative to measured signal depends on the echo time.

**Figure 5:**
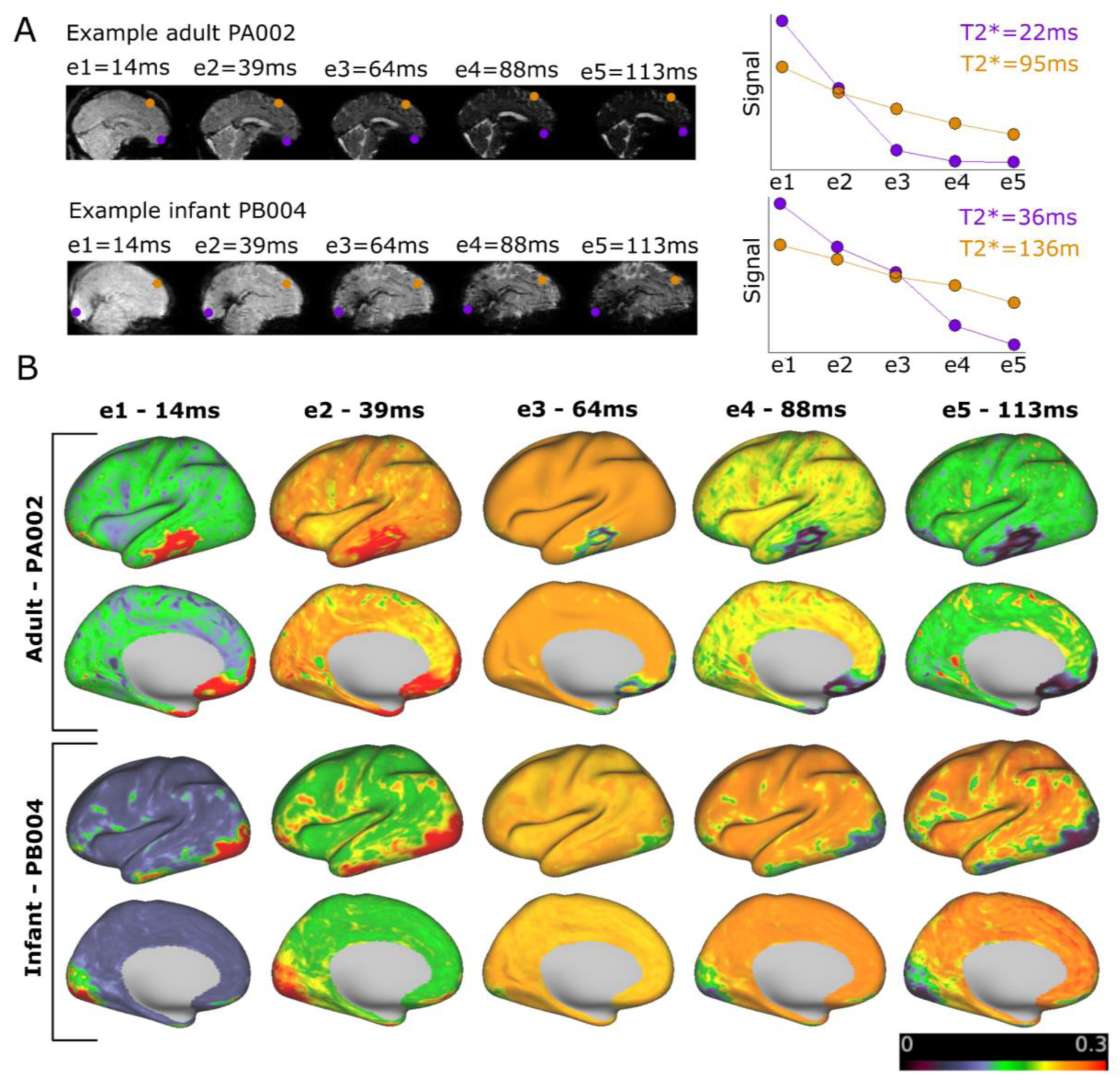
A) Example for T2 decay curve for PA002 and PB004 at two voxels with short and long T2* over five echoes. B) Echo weighting distribution across the cortex for all five echoes for an example adult and an example newborn resulting from the T2* maps displayed in Figure 4.

**Table 3:** T2* times in ms and normalized weights for each echo for the one adult and the three newborns acquired with the five-echo ME protocol. Highlighted in bold: echo weightings above 20%.

|  |  | **T2\*** | **e1** | **e2** | **e3** | **e4** | **e5** |
| --- | --- | --- | --- | --- | --- | --- | --- |
| PA002 | mean | 48.89 | 0.17 | **0.247** | **0.235** | 0.196 | 0.153 |
|  | median | 50.71 | 0.144 | 0.242 | 0.243 | 0.208 | 0.163 |
|  | SD | 12.23 | 0.095 | 0.026 | 0.031 | 0.042 | 0.043 |
|  | 5th pct | 21.26 | 0.115 | 0.215 | 0.189 | 0.091 | 0.041 |
|  | 95th pct | 64.75 | 0.347 | 0.31 | 0.243 | 0.228 | 0.2 |
| PB004 | mean | 93.84 | 0.101 | 0.194 | **0.232** | **0.240** | **0.233** |
|  | median | 98.58 | 0.085 | 0.182 | 0.232 | 0.251 | 0.25 |
|  | SD | 27.04 | 0.047 | 0.035 | 0.008 | 0.031 | 0.049 |
|  | 5th pct | 37.23 | 0.074 | 0.167 | 0.224 | 0.170 | 0.113 |
|  | 95th pct | 130.22 | 0.198 | 0.281 | 0.242 | 0.259 | 0.274 |
| PB005 | mean | 77.49 | 0.126 | **0.215** | **0.233** | **0.223** | **0.203** |
|  | median | 79.73 | 0.098 | 0.197 | 0.234 | 0.241 | 0.226 |
|  | SD | 30.46 | 0.067 | 0.046 | 0.012 | 0.043 | 0.064 |
|  | 5th pct | 27.43 | 0.075 | 0.170 | 0.212 | 0.12 | 0.062 |
|  | 95th pct | 124.6 | 0.287 | 0.319 | 0.243 | 0.258 | 0.271 |
| PB001 | mean | 81.51 | 0.114 | **0.207** | **0.234** | **0.230** | **0.214** |
|  | median | 83.51 | 0.095 | 0.194 | 0.235 | 0.243 | 0.232 |
|  | SD | 27.84 | 0.054 | 0.039 | 0.009 | 0.035 | 0.055 |
|  | 5th pct | 31.91 | 0.075 | 0.169 | 0.224 | 0.146 | 0.086 |
|  | 95th pct | 124.06 | 0.239 | 0.302 | 0.243 | 0.258 | 0.271 |

### Competition between reliability and spatial precision

To accurately characterize individual specific networks, one needs to carefully consider the tradeoff between reliability and spatial precision. This tradeoff is illustrated in Figure 6. Reducing spatial precision, for example by parcellating data, fundamentally increases reliability (Figure 6A, dotted lines vs. filled lines). However, averaging across parcels eliminates the opportunity to uncover precise individual specific patterns (Figure 6B). Thus reliability depends on spatial precision. This highlights the challenge of interpreting absolute reliability values when searching for the ideal amount of scan time. Furthermore, it is important to note that absolute reliability values also depend on the amount of held out data used to calculate them (Suppl. Fig 12).

**Figure 6:**
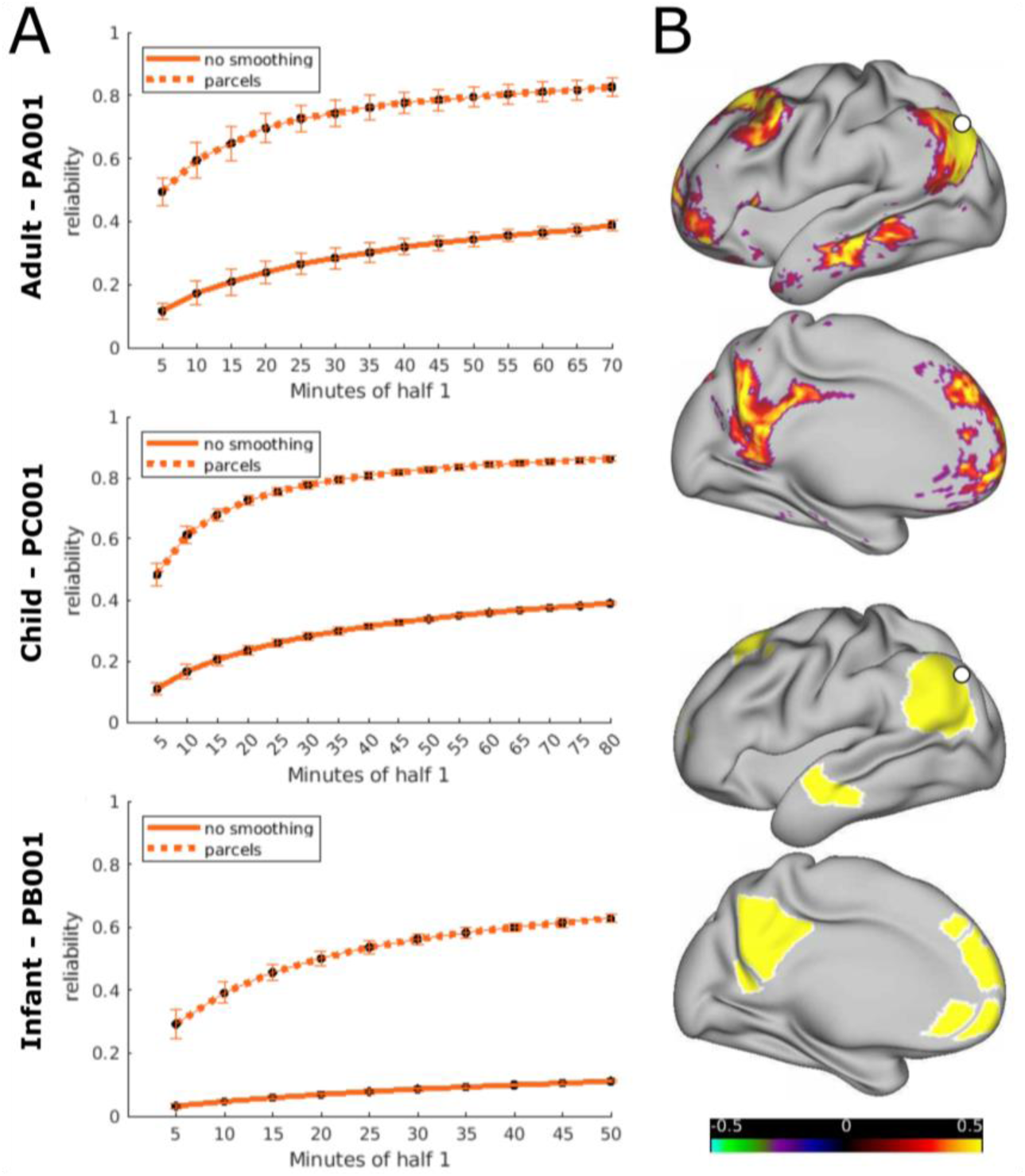
A) differences in reliability between dense, unsmoothed and parcellated ME data without NORDIC (Gordon parcels for PA001 and PC001 and infant-specific parcels for PB001). Curves represent the average reliability across all grayordinates or parcels and permutations. Error bars show SD across permutations. B) example for difference in precision between dense unsmoothed and parcellated data (ME data PA001, without NORDIC).

## Discussion

In this study, we evaluated potential methodological avenues to improve data quality and reliability to facilitate precision functional imaging, with a specific focus on developmental populations. The two methods evaluated were ME data acquisitions and NORDIC thermal denoising. In adult precision imaging subjects, we replicated prior findings showing improved tSNR and reliability with ME acquisitions (Lynch et al., 2020). Improvements in tSNR with NORDIC denoising have been previously demonstrated for SE fMRI data (Dowdle et al., 2023; Vizioli et al., 2021). Here, we demonstrated the impact of NORDIC on reliability of resting state functional connectivity and the unique benefit of combining ME and NORDIC for precision functional imaging. Expanding these findings to children and infants, we replicated the benefits of NORDIC on ME data. We furthermore used the unique feature of ME acquisitions to evaluate T2* relaxation times in all included age groups. T2* values calculated from ME data were longer and more variable in infants compared to adults. This led to a differential optimal echo combination, with heavier weighting of later echoes for the infant datasets. The ability of ME data to capture data from TEs that are closer to the infant’s T2* could be of great advantage for infant fMRI. Yet, the impact of this differential echo weighting on optimally combined ME data needs further investigation. Lastly, we demonstrated some caveats of measuring data reliability, namely its dependence on spatial precision and the available held-out data.

### PFM is feasible across development with developmentally-specific consideration

The datasets used to investigate the impact of methodological advances in the present study show that extended fMRI data acquisitions are feasible in children and even in newborn infants. For infants, collecting data over consecutive days to minimize brain developmental changes within the data acquisition period is possible. This mitigates one major constraint for precision imaging in infants, i.e., that the brain develops rapidly in early infancy (e.g., Kostović et al., (2019)). Thus, sessions for data collection need to be spaced as closely as possible in order to concatenate data without introducing variability caused by developmental changes in the brain. Even though the overall process itself was feasible, we found that motion, for example during active sleep phases or induced by spontaneous waking, is a greater challenge for this age group, than for adults or 10 year old children. Further, motion can vary substantially between individual infants, as can be seen by the difference between subjects in the amount of retained data between the subjects (PB004: ME: 82%,SE, 95%; PB005: ME: 67%, SE: 67%; PB001: 83%; Suppl. Table 1).

### NORDIC thermal noise reduction improves data quality

Our results showed an overall benefit in data quality when using NORDIC denoising in the adult, child, and infant brains. We saw an increase in tSNR as well as in the strength of functional connections and split half reliability with NORDIC. Compared to using spatial smoothing as a means to remove noise in the data and increase reliability, the application of NORDIC is preferable in order to retain spatial precision in the data (Dowdle et al., 2023; Vizioli et al., 2021). This is particularly important for infants because of their smaller brain size.

The present results suggest that NORDIC could be a valuable addition to future studies, especially since it does not come at significant cost (other than increased data storage), as it only additionally requires saving the phase data (along with the standard magnitude images). Both the acquisition of additional noise frames at the end of the scan and the usage of the default parameters (as used for PB001) proved to be beneficial. The default parameters are generally more conservative (i.e. resulting in the removal of less principal components) than the estimated threshold when using the noise frames, which allow for a 5-10% higher threshold based on a more direct estimate of thermal noise. While little effect of using the threshold based on the additional noise frames has been found in 7T auditory fMRI (Faes et al., 2024), 3T imaging might benefit from these higher thresholds (Knudsen et al., 2023).

### Advantages of ME acquisitions

ME fMRI showed improvements in data quality and reliability compared to SE acquisitions, in all subjects and across all age groups.

The evaluation of ME data acquisitions in newborn infants confirmed the findings of previous studies that showed overall longer T2* relaxation times in newborns compared to adults (Rivkin et al., 2004; Williams et al., 2005). It should be noted that T2* times vary with magnetic field strength (Peters et al., 2007) and many past infant studies used a 1.5T magnet (e.g., Leppert et al., (2009); Rivkin et al., (2004)). In addition to overall longer T2* relaxation times, our evaluation shows that newborn brains exhibit greater spatial variability of T2* compared to adults, which may be in part due to the combination of longer T2* times in developing tissues and short T2* times in areas impacted by high magnetic susceptibility effects (e.g., air – tissue interfaces). This result highlights the challenge of finding an optimal echo time for data acquisition within a given developmental age group (e.g., newborns) and supports the idea that ME fMRI could be a useful tool for imaging during brain development. Developmental studies (particularly longitudinal studies) could particularly benefit from using ME as it allows to use T2* based optimal echo weighting to dynamically adapt to developmental changes. Furthermore, the ability to estimate T2* from ME acquisitions opens up the possibility to measure other tissue properties that may be especially salient to developmental, for example, brain iron, which can be quantified by 1/T2* (R2*; e.g., Larsen et al., (2020)).

However, the impact of increased weighting of data from later echoes, which, as a function of the T2 decay curve, have weaker signals, needs to be carefully investigated. Even though BOLD activation can be detected across a wide range of TEs, results vary in sensitivity (Posse et al., 1999). Further investigations using a task fMRI design could help to better understand the use of ME acquisitions in infants. For example (Goksan et al., 2017) acquired data in infants experiencing noxious stimuli at the heel with five different SE protocols in order to identify the echo time with the maximal task contrast. Using a task design with a ME acquisition in infants could help to determine the range of echo times that optimally characterize task activation and which echoes potentially just introduce more noise. Our results, which show a large variability in T2* relaxation times in the newborn brain, suggest that it is unlikely that there is a single optimal echo time for infant fMRI. In any case, the number and timing of echoes in an ME fMRI protocol for developmental studies can still be further optimized. It will be important to consider the balance between the number of echoes, the spatial and temporal resolutions, and the acceleration factors, which are all interdependent with respect to the overall functional contrast to noise ratio. However, across all populations, our findings demonstrate the usefulness of including a relatively short TE to capture signals in areas with high signal dropout (Figure 2C).

### Advancing PFM by combining ME and NORDIC

Findings from the present study highlight the unique advantage of combining ME and NORDIC for PFM, maximally increasing reliability without sacrificing spatial precision, exactly fitting to the needs of PFM and potentially leading to improvements in other fMRI study designs as well. The large improvements of tSNR in subcortical regions are particularly promising for studies of systems that span subcortical to cortical areas and their functional development. Relative gains in tSNR and reliability with NORDIC, however, vary between individuals and between SE and ME data. One factor that needs to be considered in this context is the noise introduced by motion. The potential increase in tSNR by removal of thermal noise is limited for data in which tSNR is already compromised by motion. The infant data provide the most compelling evidence for this dependency. For example, PB004 SE and PB001 had multiple runs close to 100% low motion and showed the largest gains with NORDIC compared to the other infants that had more data compromised by motion (e.g., PB005).

Another factor that may affect the impact of NORDIC on tSNR is related to subject differences in the weighting of each echo during optimal combination of ME data. NORDIC is applied to data from each echo separately before optimally combining the data. Due to the signal decay associated with longer TEs, the relative contribution of thermal noise is different for each echo. Thus, NORDIC may have a variable benefit on tSNR depending on the relative contribution of different TEs to the optimally-combined data. Further investigations into the interaction of NORDIC and TE and the impact on optimally combined data need to be done to fully understand the combination of ME and NORDIC. This might be particularly impactful for studying infants with ME acquisitions due to their differences in echo weighing compared to children or adults.

### Considerations for investigating reliability

The quantification of reliability in this study followed the example of (Lynch et al., 2020) who performed vertex/voxel-wise correlations between dense connectivity matrices. It should be noted that reliability curves based on the correlation of entire vectorized parcellated matrices as presented in previous literature (Gordon et al., 2017; Laumann et al., 2015) yield higher values overall, which is also reflected in our parcel-wise example in Figure 6A.

The lack of a plateau in the curves we constructed indicates that the available data were not sufficient to act as suitable ‘ground truth’ held out data. This is a major difference between our analysis choice of using split-halfs and (Lynch et al., 2020), who used the correlation of the connectivity matrix of one run to the one from all other available runs of one subject (up to 5.5h of held-out data) to evaluate reliability. More generally speaking, these limitations highlight that reliability measures depend on data quantity and spatial precision, likely among other factors (e.g., TR). Interpretation of relative reliability must consider these factors when comparing values between different acquisition and analysis conditions (see Figure 6 and Suppl. Figure 12).

However, even considering the limitations to reliability quantification discussed above, reliability curves in infants seem to be overall lower in magnitude compared to children or adults. This is in line with findings by (Moore et al., 2023) and (Sylvester et al., 2022) who suggested that larger amounts of data are needed for precision imaging in infants compared to adolescents or adults. The need for longer acquisition times in infants may be explained by the general factors challenging infant imaging, such as increased motion, smaller head size relative to voxel size, increased distance to the receiver coils, or the lack of fully optimized infant specific processing pipelines (Dubois et al., 2021; Korom et al., 2021). It may, however, be in part related to more biological factors including the variability in behavioral states in newborns during scanning (quiet sleep, active sleep, wakefulness) compared to adults who are usually awake, fixating at a crosshair. Another source of variance could be small, but real, brain developmental changes across the recording days. Given the reliability limitations in infant precision functional imaging, working at the level of infant specific parcels (Myers et al., 2024; Wang et al., 2023) or individual networks (Moore et al., 2023), could help to improve reliability at the expense of spatial precision. Another approach would be to use alternative methods to increase overall SNR, such as higher magnetic fields (Annink et al., 2020), which could mitigate partial volume effects by allowing for smaller voxel sizes more fitting to the smaller infant brains, or infant specific coils (Hughes et al., 2017; Keil et al., 2011; Lopez Rios et al., 2018).

### Limitations and outlook

Comparisons between SE and ME data presented in this study are limited by the inherent differences of the acquisition sequences used, which particularly impacts tSNR. Despite this limitation, the comparison presented here reflects a potential real-world application compared to the attempt to artificially make these two sequences more similar (e.g., by using one TE from ME or subsampling SE data).

One major advantage of ME fMRI is the possibility of using MEICA for T2* based denoising (Kundu et al., 2012, 2013; Lynch et al., 2020). This is a feature we were not able to use in the present study as thresholds commonly used for MEICA are not applicable in our infant data. This is likely due to the developmental differences in T2* and will need further investigation. Given the promise of ME acquisitions for developmental neuroimaging, the development of an infant specific MEICA version could be valuable for the field, accompanied by further investigations into the effect of MEICA on spatial precision.

Furthermore, the present investigation and findings are limited as they only explored one ME fMRI protocol. More thorough investigations using different ME protocols particularly within infant subjects, and potentially even within longitudinal studies, will exploit the advantages and disadvantages of a given combination of echoes. A longitudinal approach could characterize the precise evolution of T2* relaxation times, informing the optimization of age specific protocols. The ME protocol used in this study uses a high acceleration factor (MB=6 with IPAT=2) which represents a risk of introducing spurious spatial correlations due to "slice leakage". To prevent this, we used split-slice grappa reconstruction for signal unaliasing (vs. slice-grappa) which showed to almost entirely suppress false positive activations due to slice leakage even for accelerations this high (Todd et al., 2016).

Moreover, in the present study, we did not elaborate on the role of well-trained MR operators in enhancing reliability by optimal positioning of participants in the coil and minimizing motion during data acquisition. Development of comprehensive training materials and guidelines will mitigate some of the challenges even before the availability of adequate data processing and analysis methods. As mentioned, the limitations of the present work highlight the need for more developmental specific methods developments, in particular on the avenue towards precision functional mapping during early development.

## 4. Conclusions

The present work demonstrates the utility of combining ME acquisitions with NORDIC denoising for PFM. Translating these methodological advances to developmental populations, ME data acquisitions show high promise for infant imaging, which needs further investigation. There are still gaps in our understanding of the best techniques for developmental brain imaging, motivating the development of further age-specific methodological advances as we work towards broadly applicable and robust precision functional imaging across the lifespan.

## 5. Data and Code Availability

Data can be made available upon request, given a formal data sharing agreement is set up by the institutions involved.

Code is available from the following repository: https://github.com/DCAN-Labs/code_infant_me_nordic_paper

## 6. Author Contributions

Julia Moser: Conceptualization, Formal analysis, Methodology, Software, Visualization, Writing -Original Draft; Steven M. Nelson: Conceptualization, Formal analysis, Writing - Review & Editing; Sanju Koirala: Formal analysis, Writing - Original Draft; Thomas Madison: Formal analysis, Methodology, Software, Writing - Review & Editing; Alyssa Labonte: Conceptualization, Data Curation, Writing - Review & Editing; Cristian Morales- Carrasco: Methodology, Software; Eric Feczko: Methodology, Software; Lucille A. Moore: Methodology; Jacob T. Lundquist: Formal analysis; Kimberly B. Weldon: Data Curation; Gracie Grimsrud: Formal analysis, Data Curation; Kristina Hufnagle: Data Curation; Weli Ahmed: Formal analysis; Michael J. Myers: Data Curation; Babatunde Adeyemo: Methodology; Abraham Z. Snyder: Methodology, Writing - Review & Editing; Evan M. Gordon: Writing - Review & Editing; Nico U. F. Dosenbach: Conceptualization; Brenden Trevo-Clemmens: Writing - Review & Editing; Bart Larsen: Writing - Review & Editing; Steen Moeller: Methodology, Writing - Review & Editing; Essa Yacoub: Methodology, Writing - Review & Editing; Luca Vizioli: Methodology, Writing - Review & Editing; Kamil Uğurbil: Writing - Review & Editing; Timothy O. Laumann: Conceptualization, Methodology, Writing - Review & Editing; Chad M. Sylvester: Conceptualization, Writing - Review & Editing, Supervision; Damien A. Fair: Conceptualization, Writing - Review & Editing, Supervision.

## Acknowledgements

We thank all our participants, especially the families of our precision babies for their participation and their dedication towards research. The authors acknowledge the Minnesota Supercomputing Institute (MSI) at the University of Minnesota for providing resources that contributed to the research results reported within this paper. URL: http://www.msi.umn.edu

