## Supplementary Material for "Multi-echo Acquisition and Thermal Denoising Advances Precision Functional Imaging"

### Supplemental Methods

For anatomical references, a T1 weighted scan (PA001 & children: TR = 2.5s, TE = 2.9ms, resolution = 1 x 1 x 1 mm, flip angle = 8°, infants: TR = 2.4s, TE = 2.2ms, resolution = 0.8 x 0.8 x 0.8 mm, flip angle = 8°); and a T2 weighted scan (PA001 & children: TR = 3.2s, TE = 565ms, resolution = 1 x 1 x 1 mm, flip angle = 120°, infant: TR = 4.5s, TE = 563ms, resolution = 0.8 x 0.8 x 0.8 mm, flip angle = 120°); was acquired for all participants. PA002's anatomical references were taken from a previously acquired (now publicly available) dataset (T1w: TR = 2.5s, TE = 1.81ms, resolution = 0.8 x 0.8 x 0.8 mm, flip angle = 8°; T2w: TR = 3.2s, TE = 564ms, resolution = 0.8 x 0.8 x 0.8 mm, flip angle = 120°) (Gordon et al., 2017).

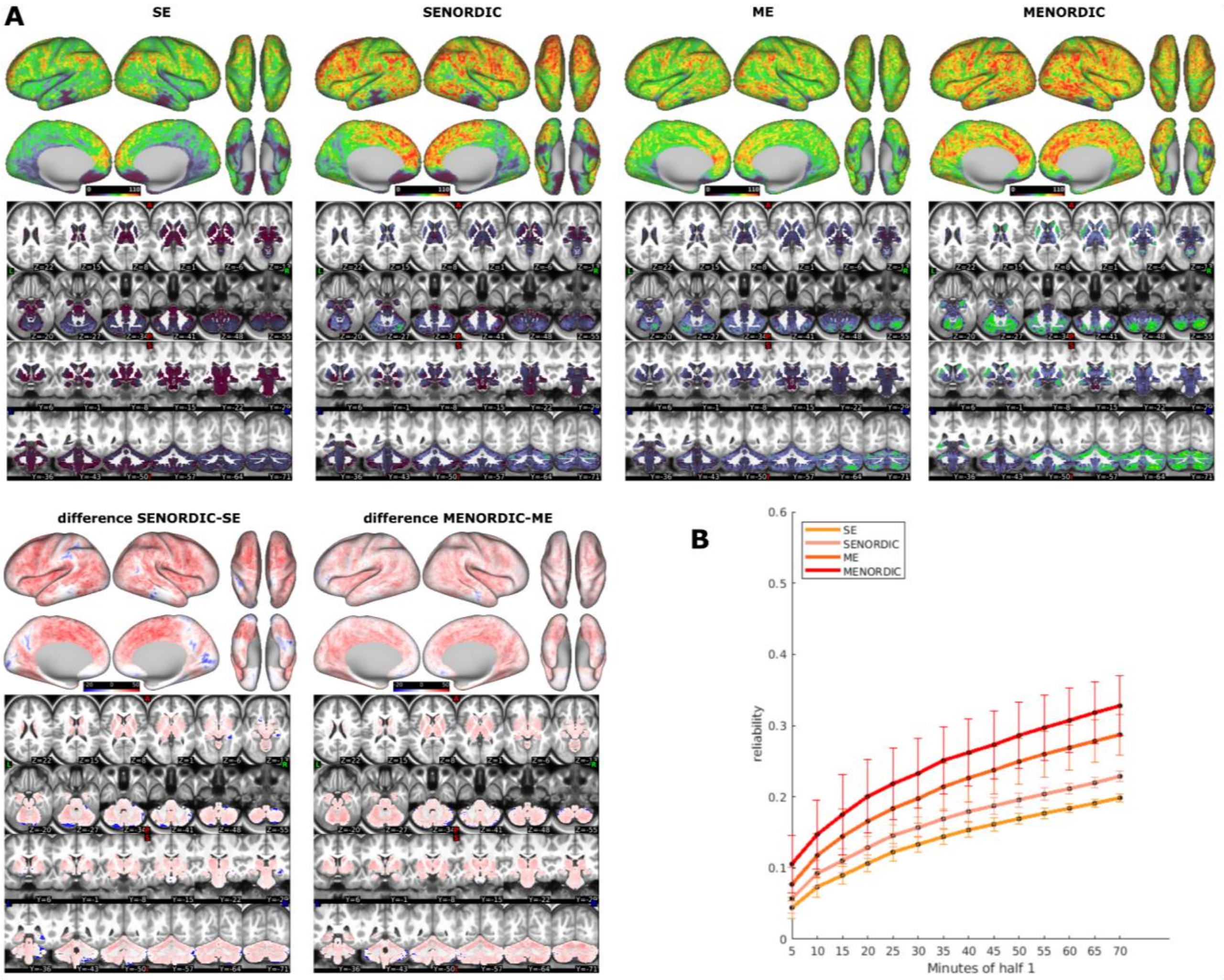


*Figure S1: A) tSNR values for cortical and subcortical structures for SE and ME data with and without NORDIC for PA002 (average of runs with >90% low motion). B) Reliability curves for a split half of the data of PA002. Curves represent the average reliability across all grayordinates and 100 permutations of the run order. Error bars show SD across permutations.*

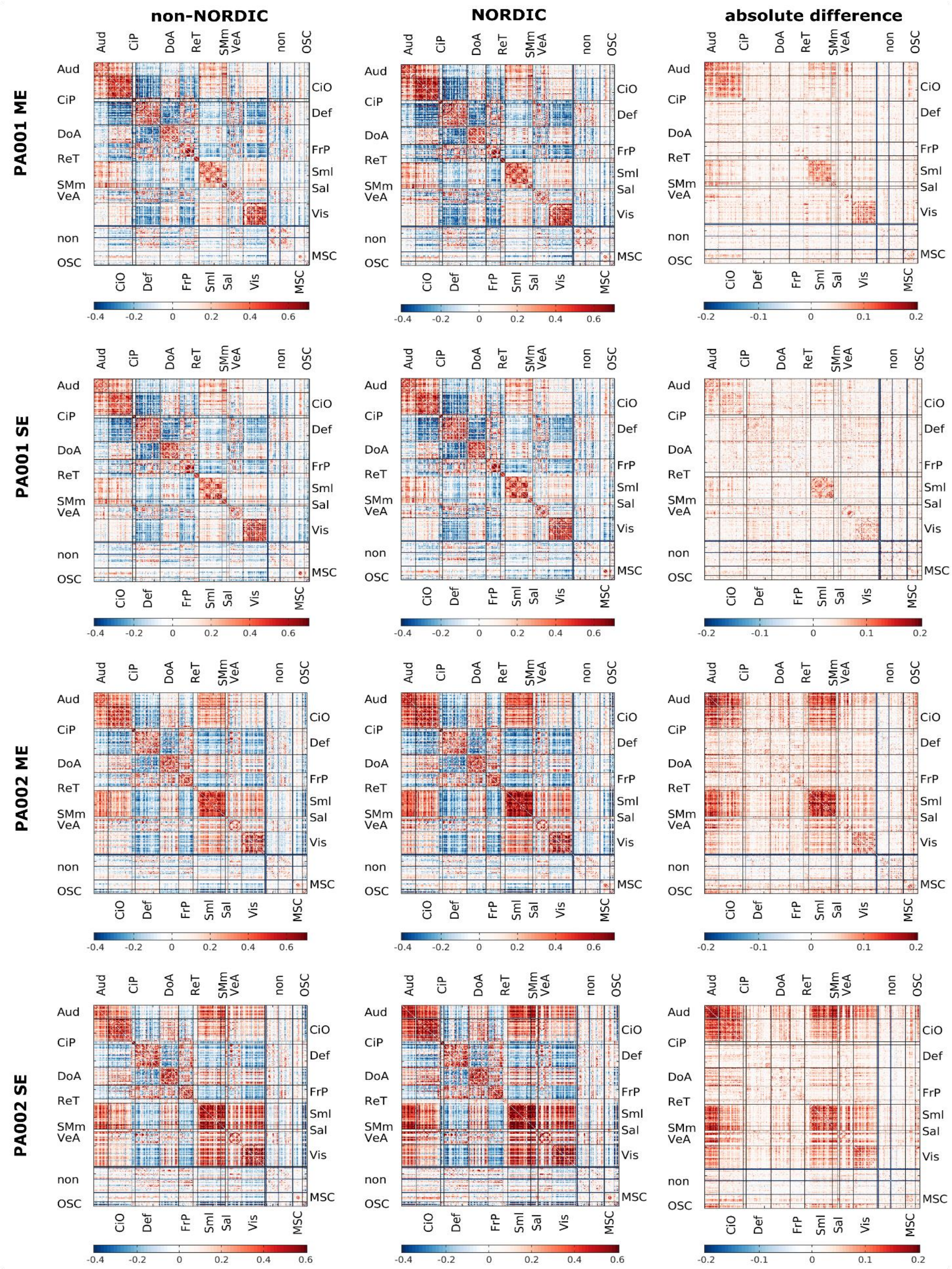


*Figure S2: Connectivity matrices from parcellated time series (Gordon parcels) showing the increase of connectivity strength with NORDIC for both ME and SE data in adults.*

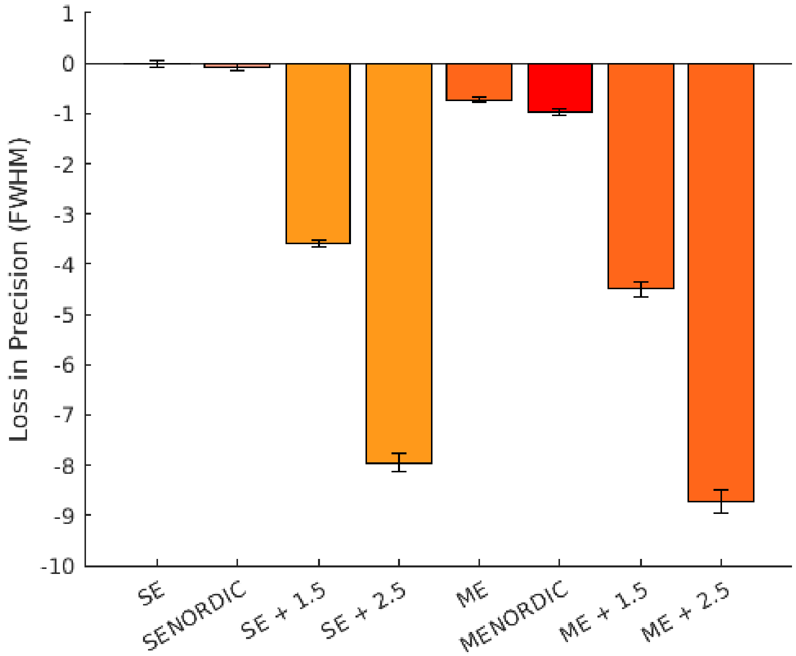


*Figure S3: loss in spatial precision with ME and NORDIC compared to commonly used smoothing kernels (σ = 1.5mm and σ =2.5mm) in example subject PA001. Spatial smoothness quantified by full width half max (FWHM) of SE data is used as reference. Error bars represent standard deviation across runs.*

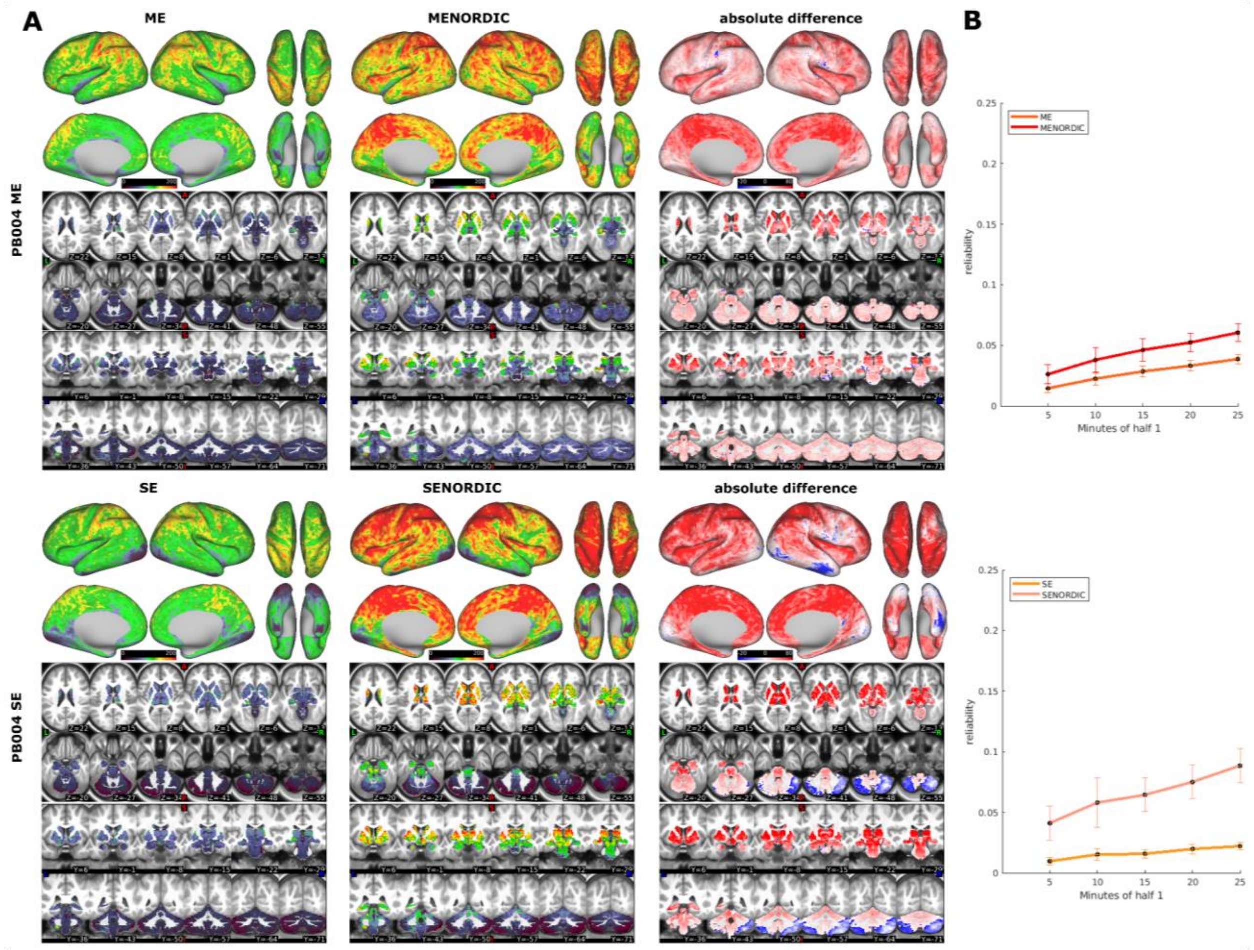


*Figure S4: increase of tSNR (A) and reliability (B) with NORDIC for ME and SE data from PB004 (tSNR represents average of runs with >90% low motion). Curves represent the average reliability across all grayordinates and 100 permutations of the run order. Error bars show SD across permutations.*

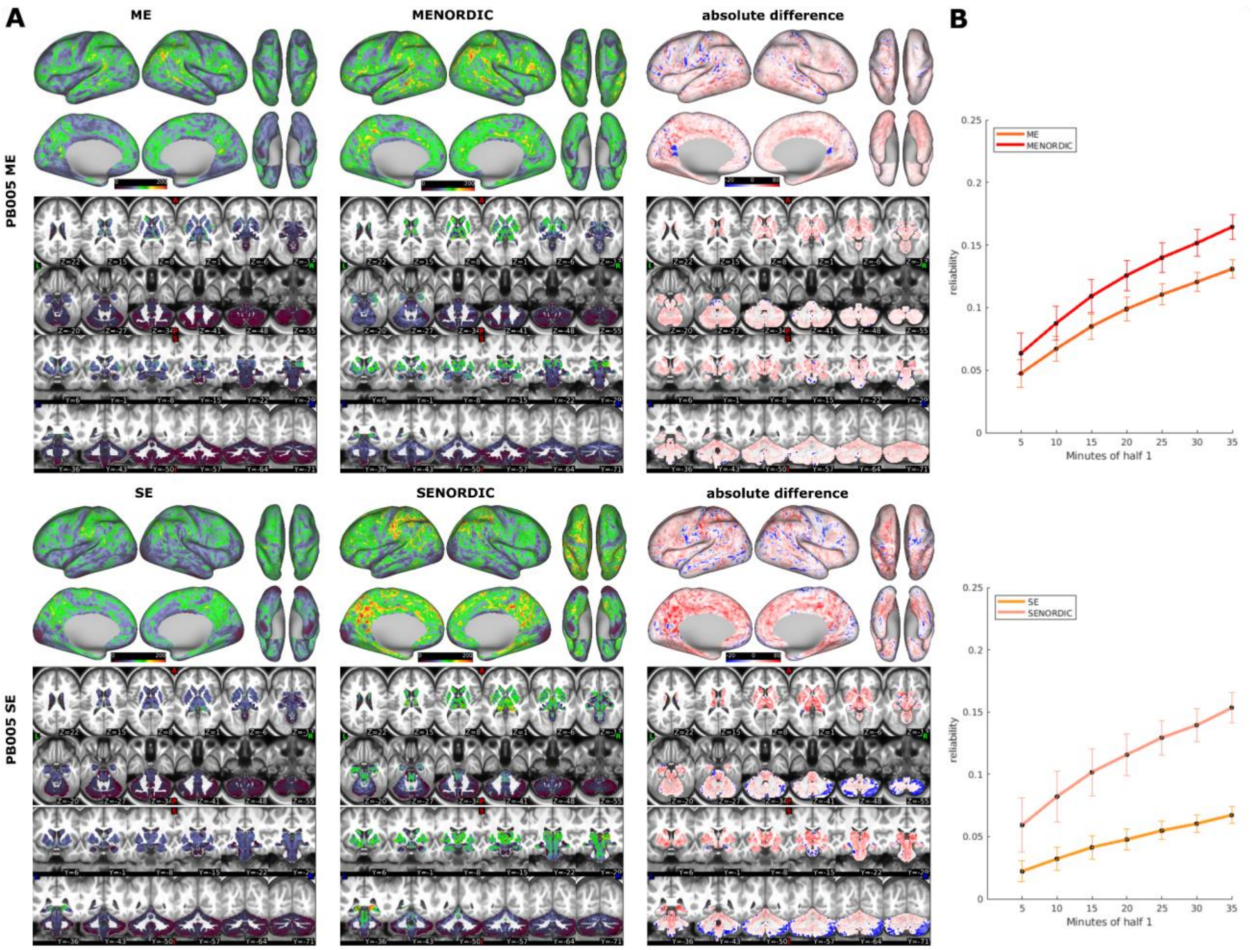


*Figure S5: increase of tSNR (A) and reliability (B) with NORDIC for ME and SE data from PB005 (tSNR represents average of runs with >90% low motion). Curves represent the average reliability across all grayordinates and 100 permutations of the run order. Error bars show SD across permutations.*

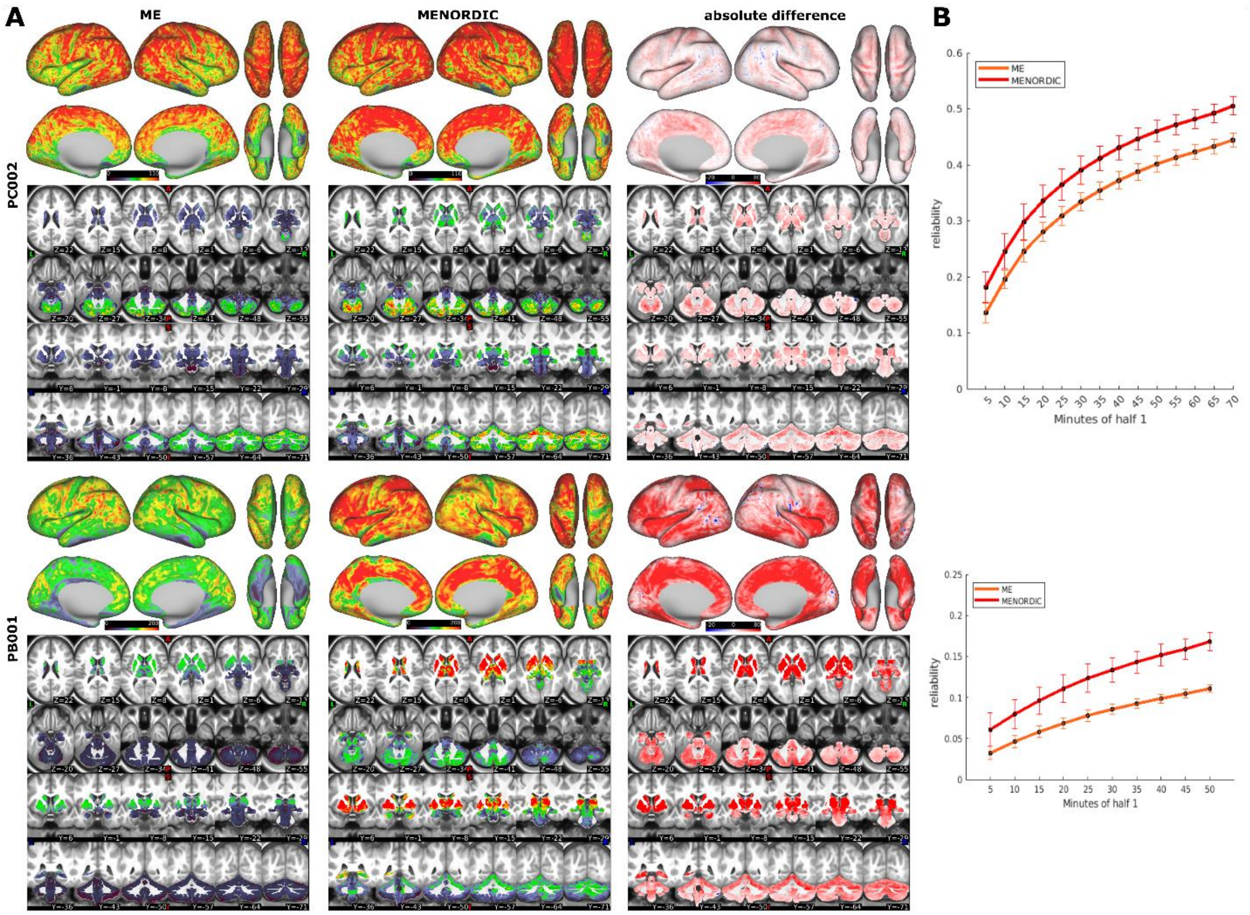


*Figure S6: increase of tSNR (A) and reliability (B) with NORDIC for ME data from PC002 and PB001 (tSNR represents average of runs with >90% low motion). Curves represent the average reliability across all grayordinates and 100 permutations of the run order. Error bars show SD across permutations.*

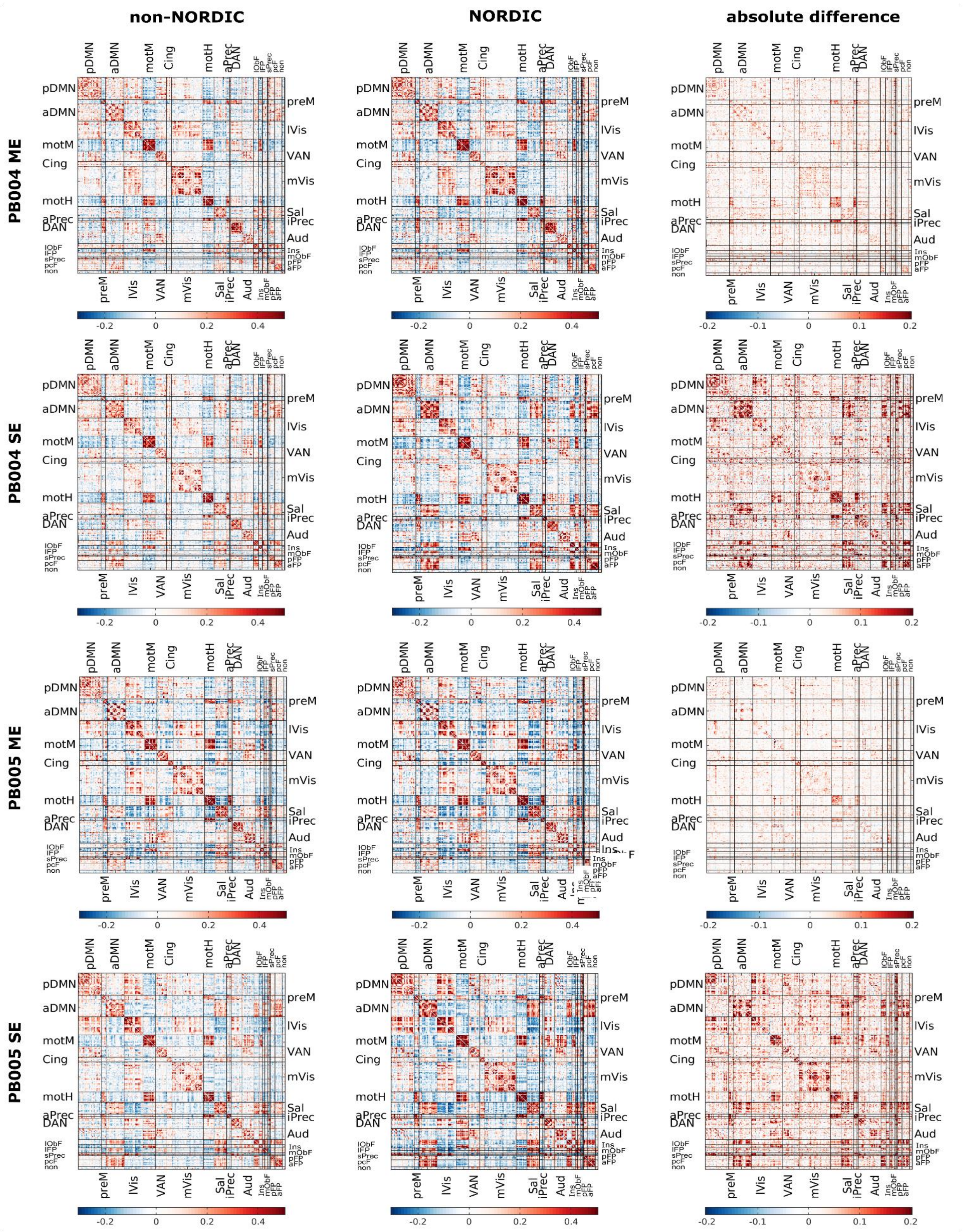


*Figure S7: Connectivity matrices from parcellated time series (baby specific parcels) showing the increase of connectivity strength with NORDIC for both ME and SE data in PB004 and PB005.*

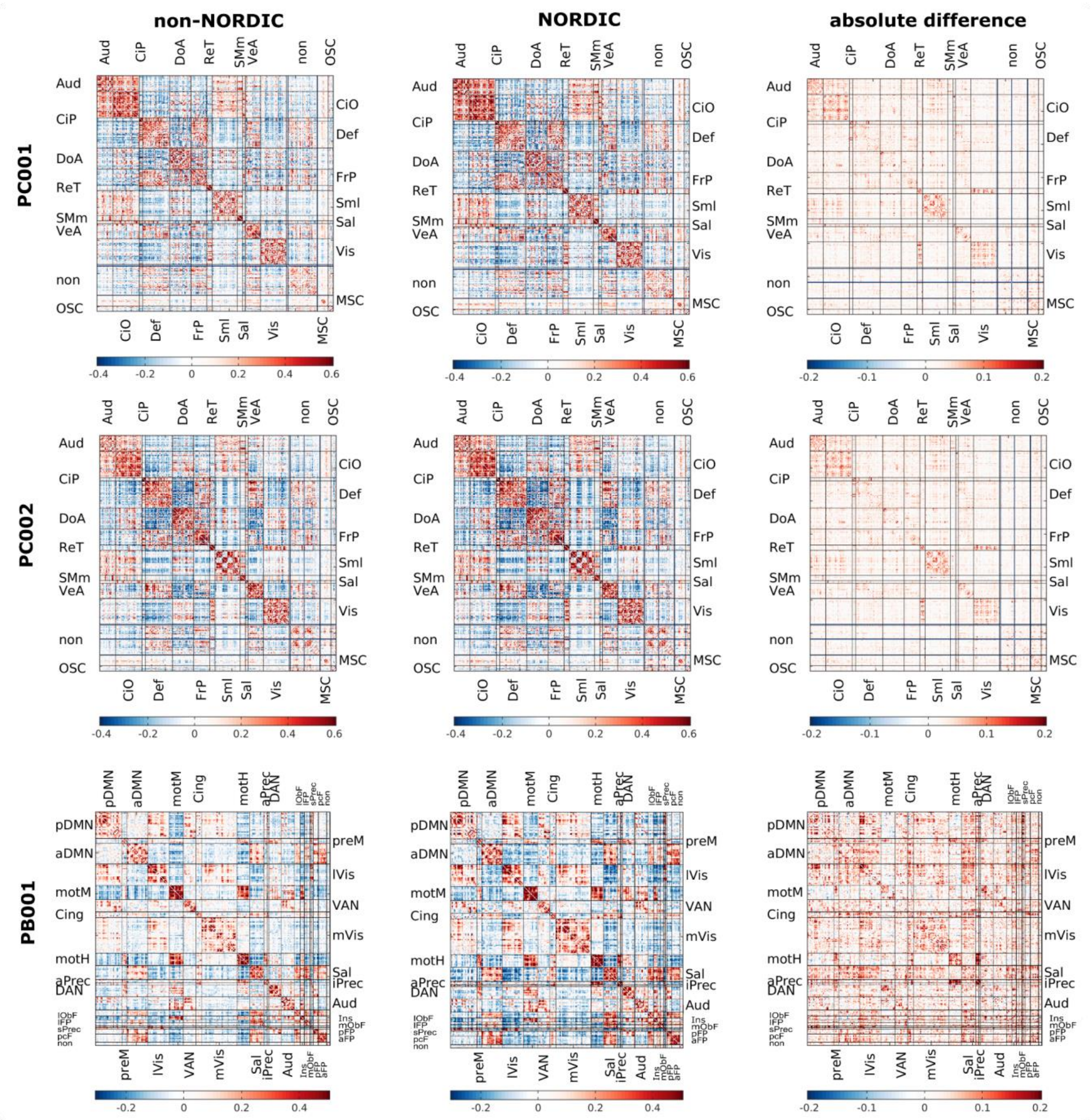


*Figure S8: Connectivity matrices from parcellated time series (Gordon parcels for PC subjects and baby specific parcels for PB subject) showing the increase of connectivity strength with NORDIC for ME data from developmental populations.*

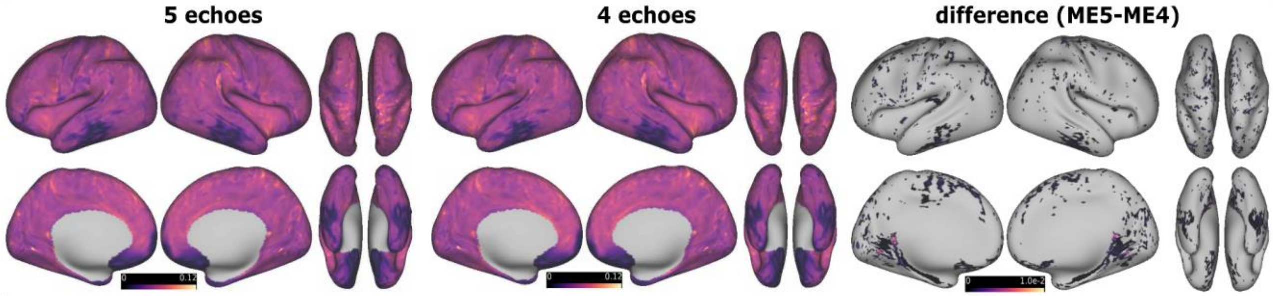


*Figure S9: T2* values (in seconds) for PA001 calculated from either all 5 echoes or the first 4 echoes, showing that using a 4 echo sequence does not impact estimation of T2* values (average of runs with >90% low motion).*

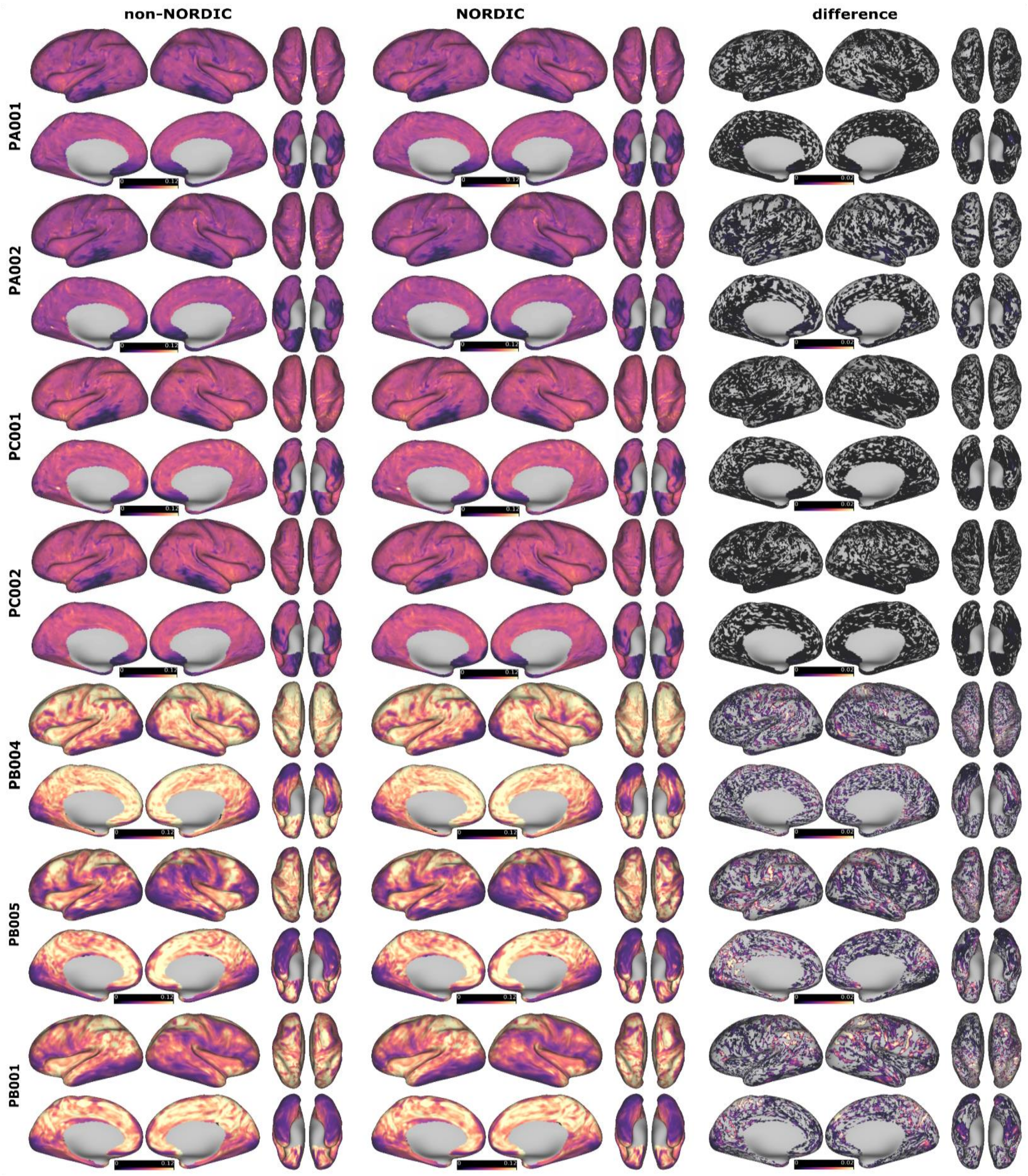


*Suppl Figure 10: T2* values (in seconds) for all precision imaging subjects for data with and without NORDIC denoising prior to echo combination (average of runs with >90% low motion). NORDIC denoising only marginally changes T2* estimates.*

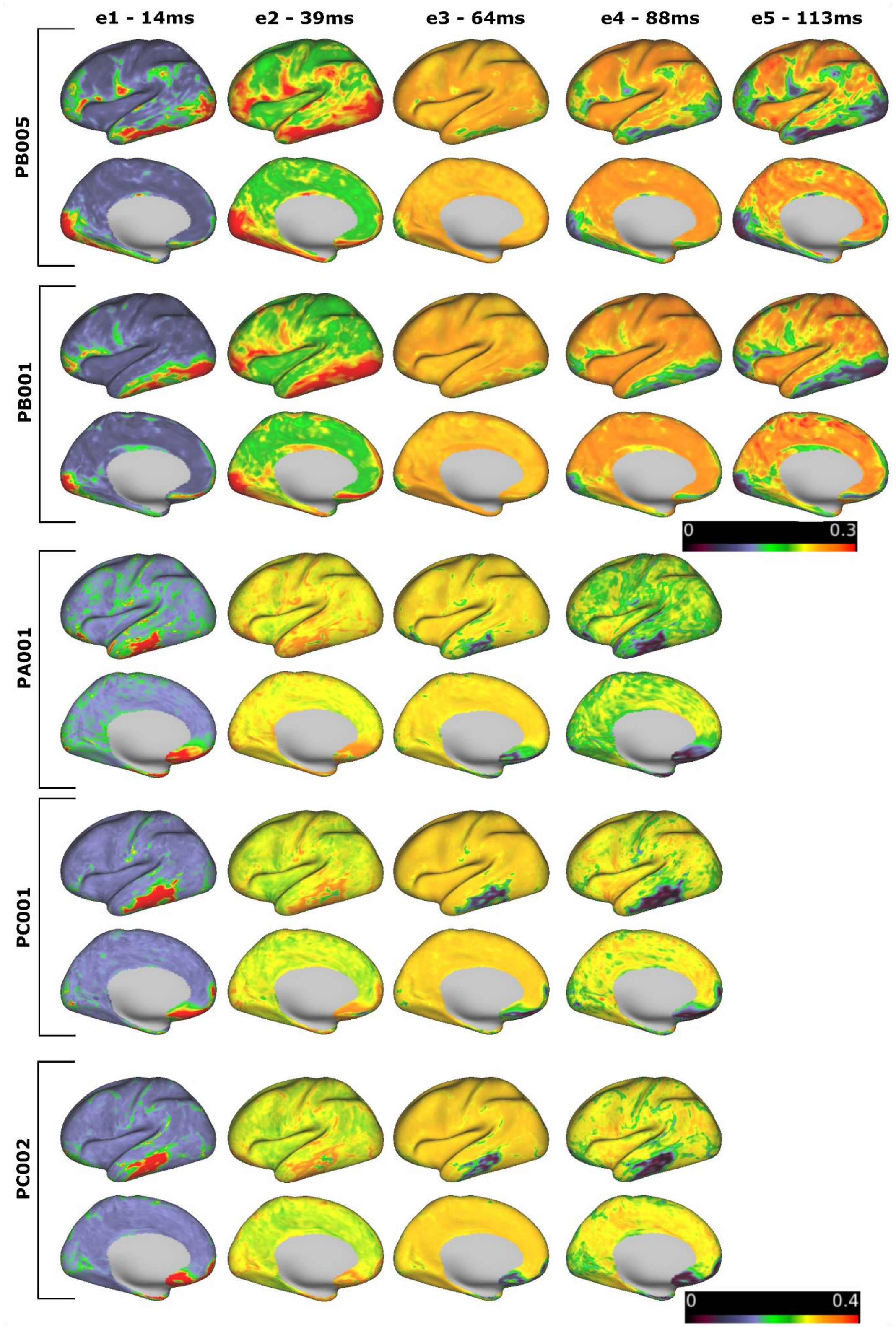


*Figure S11: T2* based echo weighting for all precision imaging subjects not depicted in Figure 5. PA001, PC001 and PC002 were acquired with a 4 echo sequence. Despite lacking the last echo, their weighting scheme very much resembles PA001 shown in Figure 5.*

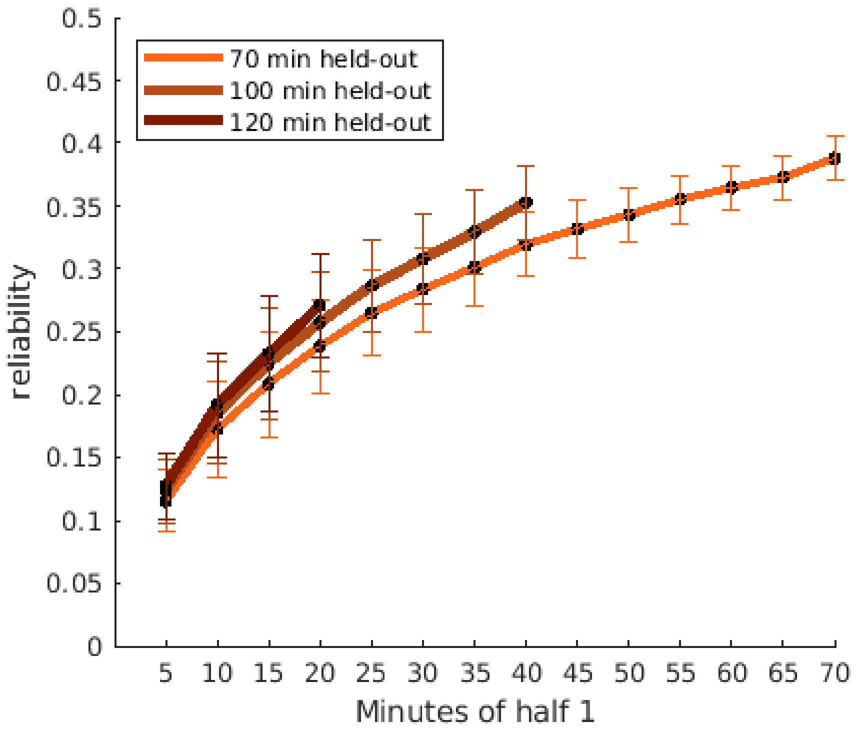


*Figure S12: Variations in reliability with variations in held out data used for calculation (ME data PA001). Curves represent the average reliability across all grayordinates and 100 permutations of the run order. Error bars show SD across permutations.*

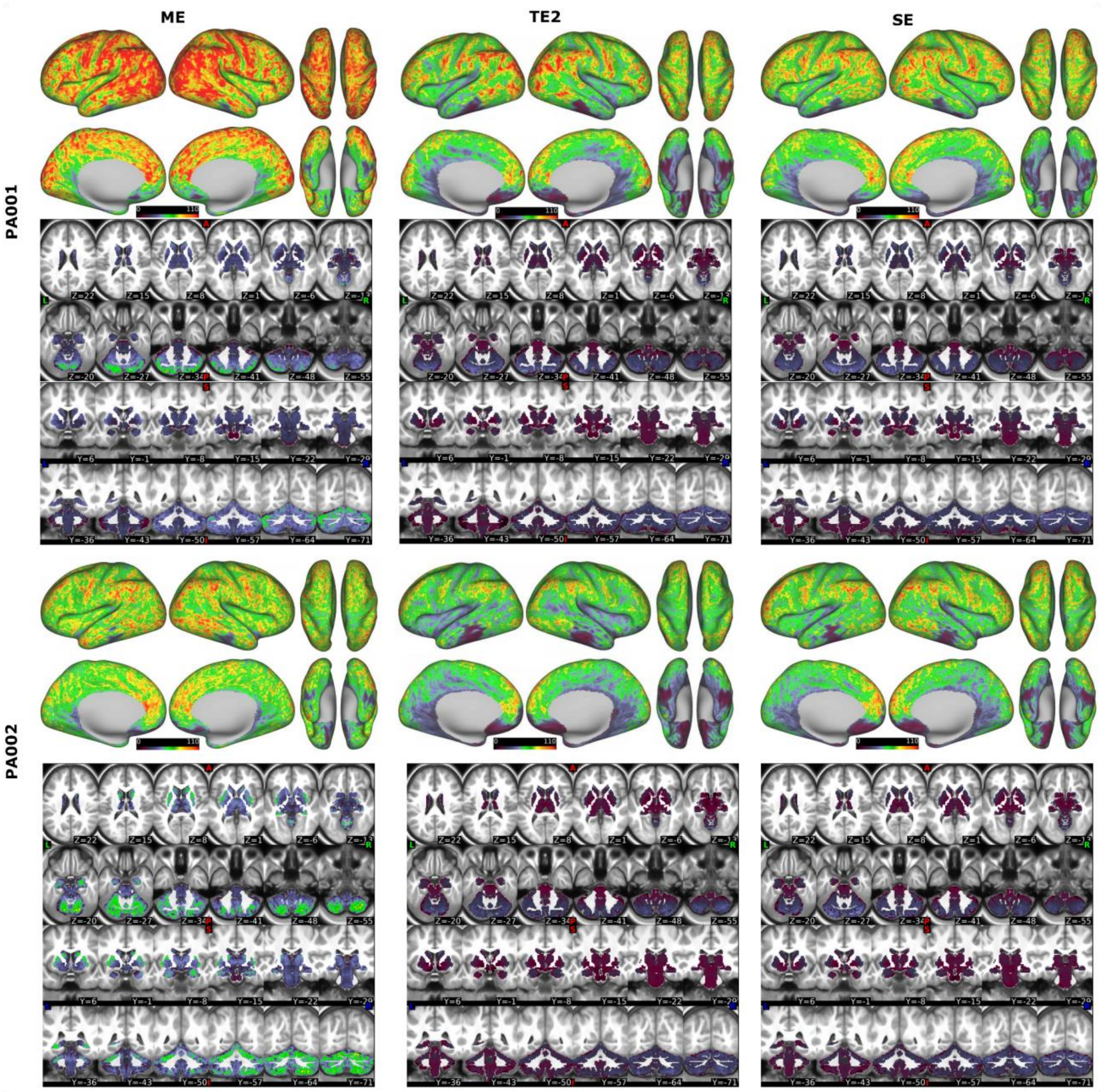


*Figure S13: tSNR comparison between ME, SE and TE2 (second echo of ME data only). This comparison addresses whether the differences in TR or flip-angle between ME and SE account for some of the improvements in tSNR seen for ME. PA001 tSNR for ME is highest (M = 67.93, SD = 31.11) compared to TE2 (M = 47.55. DS = 27.26) or SE (M = 49.23, SD = 26.13). PA002 tSNR for ME is highest (M = 54.26 , SD = 22.09) compared to TE2 (M = 40.78, SD = 22.39) or SE (M = 44.13, SD = 24.25). TE2 shows no improvements over SE.*

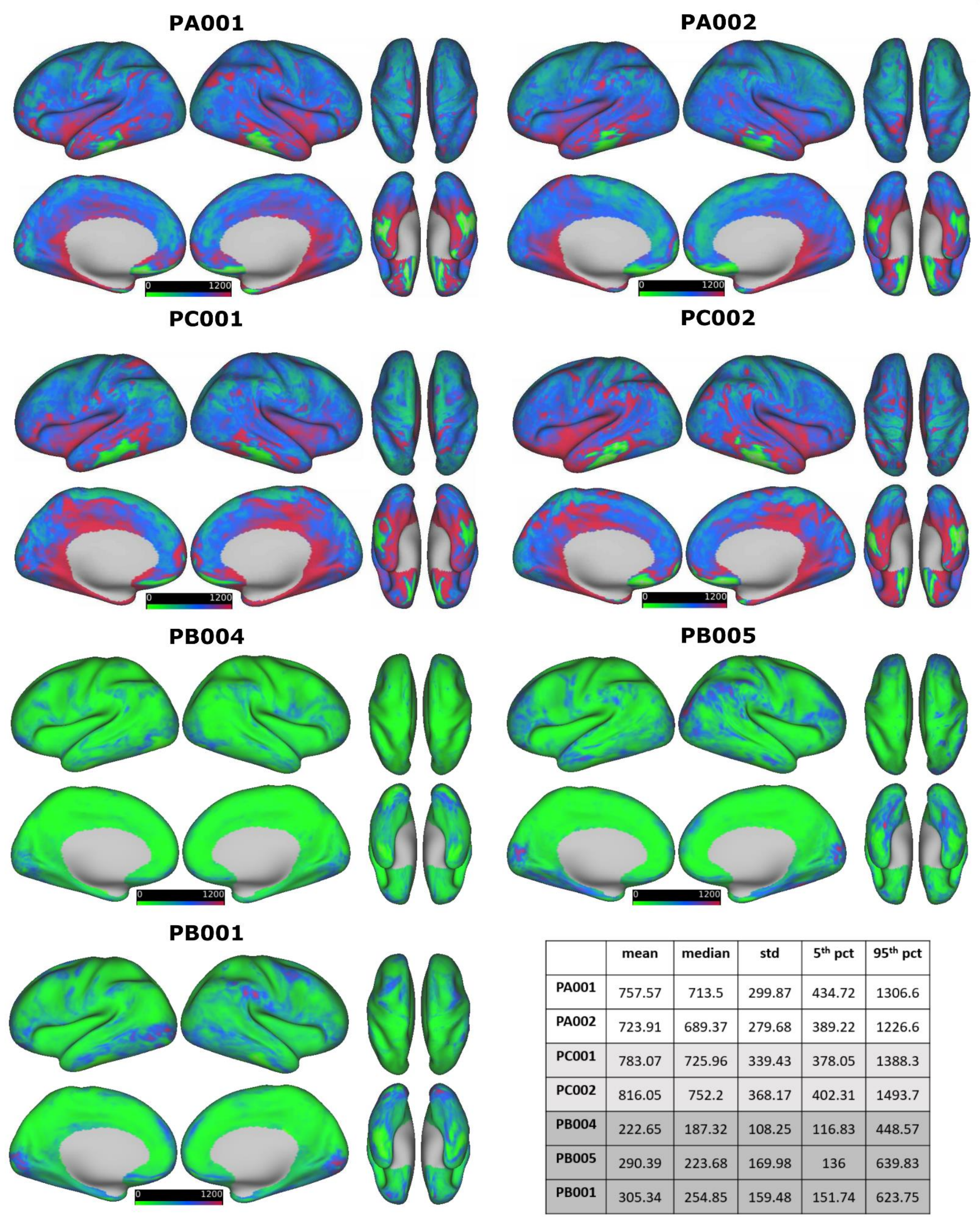


| | mean | median | std | 5th pct | 95th pct |
|---|---|---|---|---|---|
| PA001 | 757.57 | 713.5 | 299.87 | 434.72 | 1306.6 |
| PA002 | 723.91 | 689.37 | 279.68 | 389.22 | 1226.6 |
| PC001 | 783.07 | 725.96 | 339.43 | 378.05 | 1388.3 |
| PC002 | 816.05 | 752.2 | 368.17 | 402.31 | 1493.7 |
| PB004 | 222.65 | 187.32 | 108.25 | 116.83 | 448.57 |
| PB005 | 290.39 | 223.68 | 169.98 | 136 | 639.83 |
| PB001 | 305.34 | 254.85 | 159.48 | 151.74 | 623.75 |

*Figure S14: T2* and S0 model fit quantified from the root mean squared error (RMSE) output by Tedana version 24.0.1 and projected to surface space. PB subjects show smaller RMSE values compared to PC and PA subjects. It should be noted however that RMSE scales with the magnitude of the data to which the model is fit. Therefore, the values between datasets are not directly comparable.*

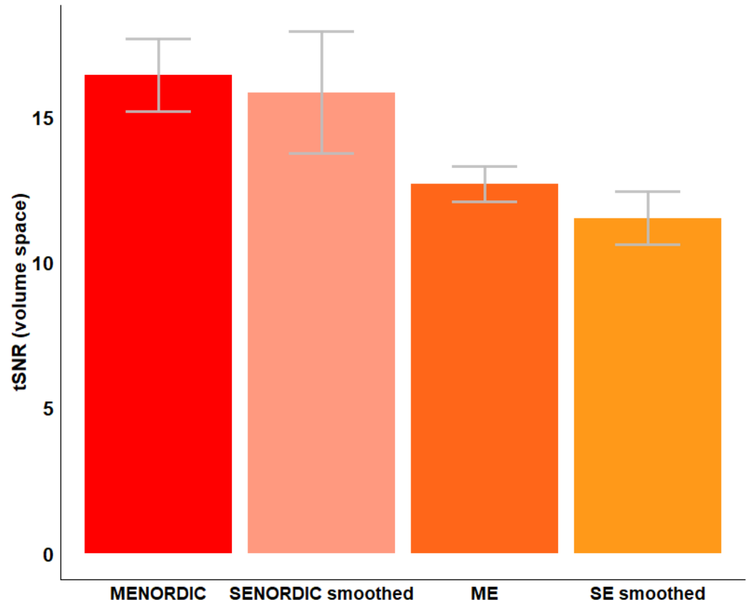


*Figure S15: Experiment on the impact of smoothing on tSNR results. Average tSNR across runs for PA001 calculated in volume space using FSL tools. Error bars represent variation across runs. SE and SE-NORDIC data are smoothed to approximately the same FWHM as ME and ME-NORDIC data using 3dBlurToFWHM in AFNI (Average FWHM for ME=3.47, SE smoothed=3.51, SE-NORDIC smoothed = 3.59, ME-NORDIC=3.7). Additional smoothing increases similarity between SE and ME data in terms of tSNR, however the relative overall pattern shown in Figure 1 (in CIFTI space) remains.*

| | **ME** | | | | **SE** | | | |
|---|---|---|---|---|---|---|---|---|
| | **Run** | **#frames** | **#good** | **percent** | **Run** | **#frames** | **#good** | **percent** |
| PA001 | 1 | 547 | 547 | 100 | 1 | 1124 | 1106 | 98.4 |
| | 2 | 547 | 547 | 100 | 2 | 1124 | 1103 | 98.13 |
| | 3 | 547 | 545 | 99.63 | 3 | 647 | 595 | 91.96 |
| | 4 | 547 | 545 | 99.63 | 4 | 1124 | 1090 | 96.98 |
| | 5 | 547 | 545 | 99.63 | 5 | 1124 | 1078 | 95.91 |
| | 6 | 547 | 546 | 99.82 | 6 | 1124 | 996 | 88.61 |
| | 7 | 547 | 541 | 98.9 | 7 | 1124 | 1041 | 92.62 |
| | 8 | 547 | 546 | 99.82 | 8 | 1124 | 1051 | 93.51 |
| | 9 | 547 | 547 | 100 | 9 | 597 | 493 | 82.58 |
| | 10 | 547 | 543 | 99.27 | 10 | 1124 | 1043 | 92.79 |
| | | | | | 11 | 597 | 495 | 82.91 |
| | | | | | 12 | 1124 | 1079 | 96 |

| | | | | | | | | |
|---|---|---|---|---|---|---|---|---|
| | | **160.54 min** | **160.02 min** | **M=99.97** | | **159.43 min** | **148.93 min** | **M=92.53** |
| PA002 | 1 | 510 | 506 | 99.22 | 1 | 1124 | 1111 | 98.84 |
| | 2 | 510 | 508 | 99.61 | 2 | 1124 | 1109 | 98.67 |
| | 3 | 441 | 416 | 94.33 | 3 | 1124 | 1115 | 99.2 |
| | 4 | 441 | 426 | 96.6 | 4 | 1124 | 1110 | 98.75 |
| | 5 | 441 | 430 | 97.51 | 5 | 1124 | 1107 | 98.49 |
| | 6 | 510 | 510 | 100 | 6 | 1124 | 582 | 51.78 |
| | 7 | 510 | 508 | 99.61 | 7 | 1124 | 1101 | 97.95 |
| | 8 | 510 | 510 | 100 | 8 | 1124 | 1097 | 97.6 |
| | 9 | 510 | 509 | 99.8 | 9 | 1124 | 1115 | 99.2 |
| | 10 | 510 | 506 | 99.22 | 10 | 1124 | 1098 | 97.69 |
| | 11 | 510 | 510 | 100 | | | | |
| | 12 | 510 | 510 | 100 | | | | |
| | 13 | 510 | 510 | 100 | | | | |
| | 14 | 510 | 510 | 100 | | | | |
| | | **203.48 min** | **201.6 min** | **M=98.99** | | **149.87 min** | **140.6 min** | **M=93.82** |
| PC001 | 1 | 547 | 547 | 100 | | | | |
| | 2 | 547 | 547 | 100 | | | | |
| | 3 | 337 | 332 | 98.52 | | | | |
| | 4 | 547 | 541 | 98.9 | | | | |
| | 5 | 547 | 546 | 99.82 | | | | |
| | 6 | 337 | 335 | 99.41 | | | | |
| | 7 | 547 | 547 | 100 | | | | |
| | 8 | 547 | 527 | 96.34 | | | | |
| | 9 | 337 | 317 | 94.06 | | | | |
| | 10 | 547 | 538 | 98.35 | | | | |
| | 11 | 547 | 545 | 99.63 | | | | |
| | 12 | 337 | 337 | 100 | | | | |
| | | **168 min** | **166.1 min** | **M=98.75** | | | | |
| PC002 | 1 | 547 | 541 | 98.9 | | | | |
| | 2 | 547 | 544 | 99.45 | | | | |
| | 3 | 337 | 337 | 100 | | | | |
| | 4 | 547 | 541 | 98.9 | | | | |
| | 5 | 547 | 546 | 99.82 | | | | |
| | 6 | 337 | 335 | 99.41 | | | | |
| | 7 | 547 | 513 | 93.78 | | | | |
| | 8 | 337 | 326 | 96.74 | | | | |
| | 9 | 547 | 508 | 92.87 | | | | |
| | 10 | 547 | 536 | 97.99 | | | | |
| | 11 | 337 | 326 | 96.74 | | | | |
| | | **151.95 min** | **148.91 min** | **M=97.69** | | | | |
| PB004 | 1 | 317 | 305 | 96.21 | 1 | 420 | 412 | 98.1 |
| | 2 | 317 | 301 | 94.95 | 2 | 131 | 131 | 100 |
| | 3 | 317 | 209 | 65.93 | 3 | 4420 | 296 | 70.48 |
| | 4 | 317 | 198 | 62.46 | 4 | 420 | 401 | 95.48 |
| | 5 | 317 | 167 | 52.68 | 5 | 420 | 416 | 99.05 |

| | | | | | | | | |
|---|---|---|---|---|---|---|---|---|
| | 6 | 317 | 304 | 95.9 | 6 | 420 | 415 | 98.81 |
| | 7 | 130 | 126 | 96.92 | 7 | 420 | 413 | 98.33 |
| | 8 | 110 | 104 | 94.55 | 8 | 420 | 142 | 98.1 |
| | 9 | 317 | 293 | 75.39 | | | | |
| | | **72.17 min** | **57.32 min** | **M=81.67** | | **77.29 min** | **72.88 min** | **M=94.79** |
| PB005 | 1 | 317 | 148 | 46.69 | 1 | 420 | 380 | 90.48 |
| | 2 | 317 | 177 | 55.84 | 2 | 420 | 356 | 84.76 |
| | 3 | 181 | 134 | 74.03 | 3 | 420 | 286 | 68.1 |
| | 4 | 317 | 272 | 85.80 | 4 | 420 | 172 | 40.95 |
| | 5 | 317 | 279 | 88.01 | 5 | 284 | 94 | 33.1 |
| | 6 | 317 | 149 | 47 | 6 | 420 | 399 | 95 |
| | 7 | 317 | 162 | 51.1 | 7 | 420 | 321 | 76.43 |
| | 8 | 317 | 292 | 92.11 | 8 | 115 | 61 | 53.04 |
| | 9 | 317 | 310 | 97.79 | 9 | 301 | 198 | 65.78 |
| | 10 | 317 | 209 | 65.93 | 10 | 420 | 337 | 80.24 |
| | 11 | 317 | 173 | 54.57 | 11 | 420 | 267 | 63.57 |
| | 12 | 317 | 258 | 81.39 | 12 | 420 | 293 | 69.76 |
| | 13 | 317 | 103 | 32.49 | 13 | 420 | 227 | 54.05 |
| | | **116.96 min** | **78.25 min** | **M=67.14** | | **123.32 min** | **85.34 min** | **M=67.33** |
| PB001 | 1 | 230 | 226 | 98.26 | | | | |
| | 2 | 230 | 227 | 98.7 | | | | |
| | 3 | 230 | 135 | 58.7 | | | | |
| | 4 | 230 | 77 | 33.48 | | | | |
| | 5 | 230 | 217 | 94.35 | | | | |
| | 6 | 230 | 227 | 98.7 | | | | |
| | 7 | 230 | 110 | 47.83 | | | | |
| | 8 | 230 | 226 | 98.26 | | | | |
| | 9 | 230 | 223 | 96.96 | | | | |
| | 10 | 230 | 225 | 97.83 | | | | |
| | 11 | 230 | 196 | 85.22 | | | | |
| | 12 | 230 | 67 | 29.13 | | | | |
| | 13 | 230 | 227 | 98.7 | | | | |
| | 14 | 230 | 227 | 98.7 | | | | |
| | 15 | 230 | 226 | 98.26 | | | | |
| | 16 | 230 | 146 | 52.17 | | | | |
| | 17 | 230 | 120 | 80.87 | | | | |
| | 18 | 230 | 186 | 80.88 | | | | |
| | 19 | 230 | 207 | 90 | | | | |
| | 20 | 230 | 221 | 96.09 | | | | |
| | 21 | 230 | 170 | 73.91 | | | | |
| | | **141.76 min** | **112.09 min** | **M=83.02** | | | | |

*Supplementary Table 1: Frames per run and frames per run remaining after motion censoring. Infants: FD<0.3, adults FD<0.2; gray: excluded as below 30% of the run.*

| | amount of data | | non-NORDIC | NORDIC | absolute difference |
|---|---|---|---|---|---|
| PA001 | 140 min | ME | M =0.172, SD =0.407 | M =0.193, SD =0.411 | M =0.028, SD = 0.026 |
| | | | min=-0.765, max= 1.955 | min=-0.784, max= 1.988 | min=-0.087, max= 0.25 |
| | | SE | M =0.16, SD =0.403 | M =0.178, SD =0.407 | M =0.028, SD =0.025 |
| | | | min=-0.733, max= 1.767 | min=-0.758, max= 1.756 | min=-0.705, max= 0.255 |
| PA002 | 140 min | ME | M =0.179, SD =0.405 | M =0.211, SD = 0.413 | M =0.042, SD = 0.04 |
| | | | min=-0.69, max= 1.61 | min=-0.748, max= 1.716 | min=-0.328, max= 0.388 |
| | | SE | M =0.18, SD =0.407 | M =0.213, SD =0.416 | M =0.04, SD =0.04 |
| | | | min=-0.805, max= 1.527 | min=-0.84, max= 1.543 | min=-0.123, max= 0.307 |
| PC001 | 160 min | ME | M =0.156, SD = 0.401 | M =0.172, SD = 0.405 | M =0.022, SD = 0.022 |
| | | | min= -0.544, max= 1.575 | min=-0.569, max= 1.573 | min=-0.107, max= 0.285 |
| PC002 | 140 min | ME | M =0.161, SD =0.404 | M =0.176, SD = 0.408 | M =0.022, SD = 0.02 |
| | | | min=-0.577, max= 1.653 | min=-0.598, max= 1.684 | min=-0.1, max= 0.247 |
| PB004 | 55 min | ME | M = 0.122, SD = 0.436 | M = 0.135, SD = 0.438 | M = 0.029, SD = 0.025 |
| | | | min=-0.346, max=1.276 | min=-0.377, max=1.391 | min=-0.136, max=0.261 |
| | | SE | M =0.115, SD =0.434 | M =0.141, SD =0.441 | M =0.051, SD =0.052 |
| | | | min=-0.331, max= 1.155 | min=-0.405, max= 1.497 | min=-0.235, max= 0.581 |
| PB005 | 75 min | ME | M = 0.128, SD = 0.438 | M = 0.14, SD = 0.44 | M = 0.024, SD = 0.022 |
| | | | min=-0.455, max=1.395 | min=-0.493, max=1.45 | min=-0.197, max=0.262 |
| | | SE | M =0.12, SD =0.435 | M =0.151, SD =0.442 | M =0.049, SD =0.048 |
| | | | min=-0.352, max= 1.078 | min=-0.418, max= 1.646 | min=-0.203, max= 0.599 |
| PB001 | 110 min | ME | M =0.121, SD = 0.437 | M =0.146, SD = 0.44 | M = 0.044, SD = 0.04 |
| | | | min=-0.34, max=1.376 | min=-0.428, max=1.427 | min=-0.302, max=0.406 |

*Supplementary Table 2: overall connectivity strength of parcellated data with and without NORDIC for precision imaging participants. All fisher z-transformed values, range excludes the matrix diagonal.*

| | | non-NORDIC | non-NORDIC >90% low motion runs only | NORDIC | NORDIC >90% low motion runs only |
|---|---|---|---|---|---|
| PA001 | ME | 3.47 | 3.47 | 3.7 | 3.7 |
| | SE | 2.73 | 2.73 | 2.8 | 2.8 |
| PA002 | ME | 3.26 | 3.26 | 3.35 | 3.35 |
| | SE | 2.5 | 2.5 | 2.51 | 2.51 |
| PC001 | ME | 3.25 | 3.25 | 3.43 | 3.43 |
| PC002 | ME | 3.39 | 3.39 | 3.61 | 3.61 |
| PB004 | ME | 7.02 | 6.8 | 7.41 | 7.25 |
| | SE | 6.28 | 6.26 | 6.6 | 6.58 |
| PB005 | ME | 8.48 | 7.89 | 9.13 | 8.53 |
| | SE | 7.25 | 6.97 | 7.69 | 7.46 |
| PB001 | ME | 7.73 | 7.37 | 8.5 | 8.31 |

*Supplementary Table 3: Data smoothness estimated by full width half max (FWHM) for all participants. Table lists mean across all runs and all runs with >90% low motion data. Note that all data are in MNI space, which increases smoothing for infant data compared to children and adults.*
